# CREsted: modeling genomic and synthetic cell type-specific enhancers across tissues and species

**DOI:** 10.1101/2025.04.02.646812

**Authors:** Niklas Kempynck, Seppe De Winter, Casper H. Blaauw, Vasileios Konstantakos, Sam Dieltiens, Eren Can Ekşi, Valérie Bercier, Ibrahim I. Taskiran, Gert Hulselmans, Katina Spanier, Valerie Christiaens, Ludo Van Den Bosch, Lukas Mahieu, Stein Aerts

**Author notes:** lllumina Artificial Intelligence Laboratory, Illumina Inc.; Foster City, California, 94404, USA. These authors contributed equally to this work (Niklas Kempynck, Seppe De Winter).

## Abstract

Sequence-based deep learning models have become the state of the art for the analysis of the genomic regulatory code. Particularly for transcriptional enhancers, deep learning models excel at deciphering sequence features and grammar that underlie their spatiotemporal activity. To enable end-to-end enhancer modeling and design, we developed a software and modeling package, called CREsted. It combines preprocessing starting from single-cell ATAC-seq data; modeling with a choice of several architectures for training classification and regression models on either *topics* or pseudobulk peak heights; sequence design using multiple strategies; and downstream analysis through a collection of tools to locate transcription factor (TF) binding sites, infer the effect of a TF (activating or repressing) on enhancer accessibility, decipher enhancer grammar, and score gene loci. We demonstrate CREsted using a mouse cortex model that we validate using the BICCN collection of *in vivo* validated mouse brain enhancers. Classical enhancers in immune cells, including the *IFNB1* enhanceosome are revisited using a PBMC model, and we assess the accuracy of TF binding site predictions with ChIP-seq. Additionally, we use CREsted to compare mesenchymal-like cancer cell states between tumor types; and we investigate different fine-tuning strategies of Borzoi within CREsted, comparing their performance and explainability with CREsted models trained from scratch. Finally, we train a CREsted model on a scATAC-seq atlas of zebrafish development and use this to design and *in vivo* validate cell type-specific synthetic enhancers in three tissues. For varying datasets, we demonstrate that CREsted facilitates efficient training and analyses, enabling scrutinization of the enhancer logic and design of synthetic enhancers across tissues and species. CREsted is available at https://crested.readthedocs.io.

## Introduction

Cell type identity is encoded in the genome in non-coding *cis*-regulatory elements^1^. In particular enhancers are important elements that harbor specific combinations of transcription factor binding sites (TFBS). This combination of TFBS, for activators and repressors, their strength, copy number, and relative arrangement are referred to as the “enhancer code”. However, both TF binding and the enhancer code itself are degenerate. That is, a single TF can bind to multiple DNA sequences (TF binding degeneracy) and different combinations of a TFBS set can generate a functional enhancer for a given cell type (enhancer degeneracy)^2^. Therefore, to model enhancers, methods are needed that can handle this diversity. Deep neural networks are particularly well suited for this^3–11^. So called sequence-to-function models^12^ take in genomic sequences and predict a variety of genomic assays, including TF binding^9^, chromatin accessibility^4,6–8,13^, enhancer activity^14^ and gene expression^5,10,15^. Among those, single-cell assay for transposase-accessible chromatin using sequencing (scATAC-seq) has shown great promise in identifying cell types and their specifically active regulatory elements, revealing cellular heterogeneity and reconstructing differentiation trajectories^16–21^. Moreover, differential region accessibility across cell types has been found to be a strong indicator of cell type-specific enhancer function, in different species and biological systems^22–24^. Therefore, sequence-to-function models trained on scATAC-seq data have been prominently used to decode enhancers.

Given that the expression of most genes is regulated by multiple enhancers, some modeling approaches have focused on using large sequence contexts. These methods often predict multiple genomic assays to achieve a global view of gene regulation, including bulk or scRNA-seq^5,10,15,25,26^. These studies have mostly focused on predicting the effect of variants on gene expression, but have not been used extensively to decode enhancer logic. Other approaches have focused on local enhancer contexts, with the aim of capturing the sequence features that define their functionality at high resolution^4,6,8,11,27^. Such models have been used to describe enhancers across tissues and species, deciphering their inherent codes^13,14,21,28–30^. Additionally, with the aim of synthetically designing enhancers, these local enhancer models have been the modeling option of choice^31–33^.

However, these local enhancer models were mostly designed to address specific questions on a fixed dataset and/or by a fixed model architecture; thus, hindering their extensibility and reproducibility. To facilitate and standardize the application of deep learning in regulatory genomics, software packages have recently been introduced with the aim of streamlining the process of data processing and model training across different datasets and architectures^8,27,34,35^. Packages such as Selene^36^ and EUGENe^34^ offer frameworks for various predictive tasks and model architectures. Other methods like *ChromBPNet*^8^ and *scPrinter*^27^ focus on predicting TF footprinting to obtain insights into enhancer codes. Footprinting modeling does require a large amount of data (large number of cells with high sequencing depths ideally over 100M reads), which is not straightforward to obtain in complex tissues at single cell resolution. Additionally, both ChromBPNet and scPRINTER do not focus on sequence design implementations. Lastly, gReLU^35^ is a comprehensive toolkit covering many steps in DNA sequence modeling such as data processing, model training, variant effect prediction, and model-guided sequence design. While all these frameworks in general share some features, they can be complex to integrate into modern scATAC-seq data analysis pipelines; they have not been validated on large scale and complex scATAC-seq datasets in different biological systems; and they often lack comprehensive global enhancer code analysis tools.

Here we present CREsted (Cis Regulatory Element Sequence Training, Explanation, and Design), a Python software package integrated into the *scverse*^37^ providing user-friendly deep learning modeling of scATAC-seq data combined with a complete analysis of enhancer code at a cell type-specific, nucleotide-level resolution. By examining various biological systems, such as the mouse motor cortex, human peripheral blood mononuclear cells (PBMC), human cancer cell states, and the developing zebrafish, we demonstrate that CREsted is versatile across species and tissues, capable of handling large datasets. We validate the identification of relevant TFBS in these systems, highlight cross-species prediction capabilities and *in vivo* validate sequence design methods from our CREsted models. This work not only showcases the robustness and applicability of CREsted but also paves the way for further advancements in the field of regulatory genomics.

## Results

### CREsted is a software package for efficient enhancer modeling and design

CREsted consists of four main modules: data preprocessing, model training, model interpretation, and synthetic enhancer design (Fig. 1). It starts from processed scATAC-seq data and provides, through the analysis of genomic sequences and the design of synthetic sequences, a thorough understanding of the enhancer code of cell states or types in the biological system under study.

**Fig. 1:**
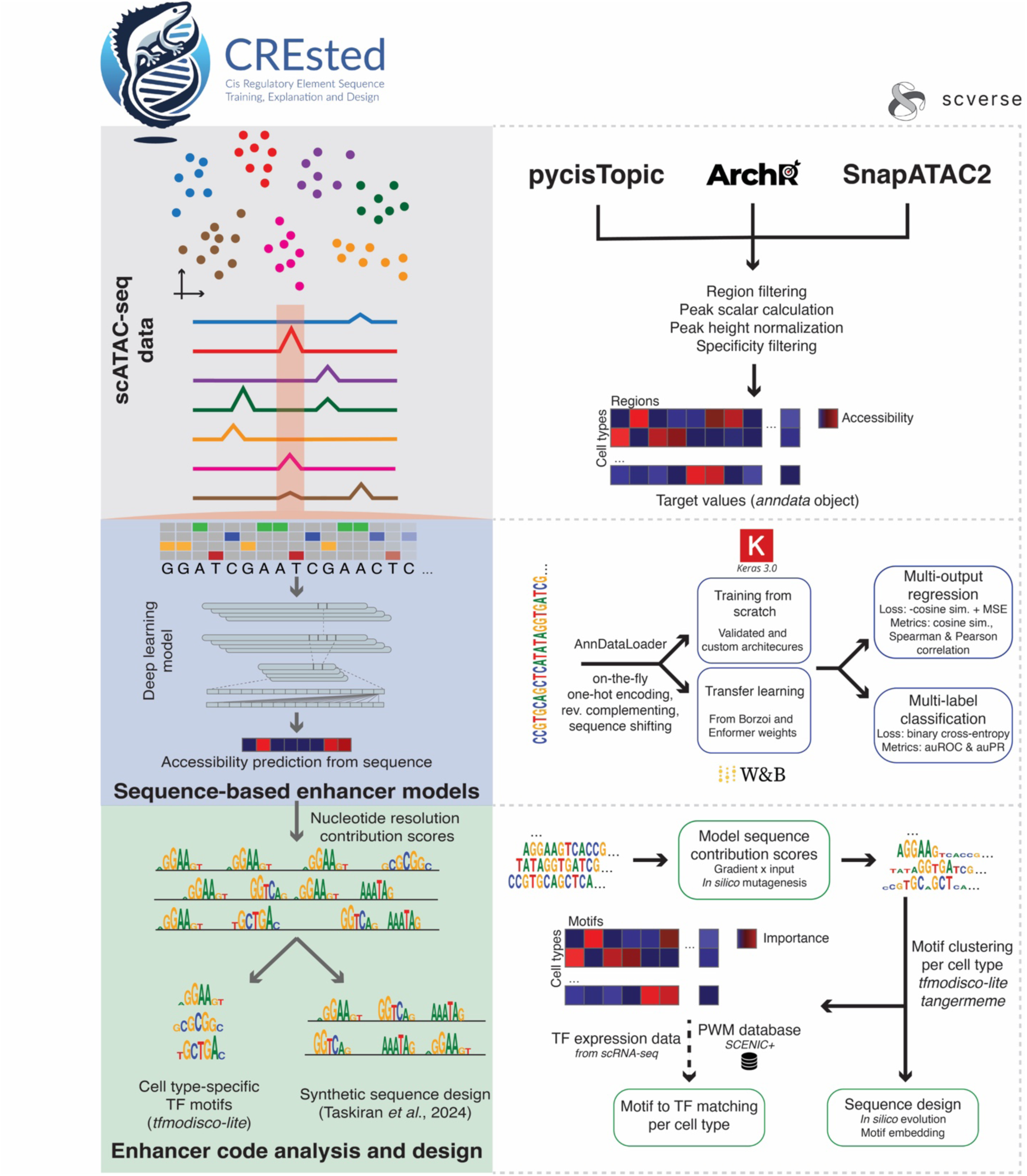
CREsted package overview. CREsted is a software package for training enhancer models, analyzing cell type-specific enhancer codes and designing synthetic enhancer sequences. From preprocessed scATAC-seq data, we apply additional steps to obtain accessibility values per cell type. Those are used as target values to train sequence-based enhancer models that can be trained from scratch or transfer learned from large-scale models. Both multi-output regression and multi-label classification are possible. From the trained models, we obtain insights into cell type-specific enhancer codes by looking at nucleotide contribution scores to identify motifs and match them to TF candidates. Finally, CREsted allows for synthetic sequence design using the obtained enhancer code insights.

Data preprocessing builds on the output of established scATAC-seq data analysis pipelines^38–40^ and has two modes. The first mode is based on topic modeling from pycisTopic^39,41^, whereby cells and peaks are assigned to latent topics representing matched probability distributions over peaks and single cells. The second mode uses pseudobulk peak heights, represented by counts per million (CPM) normalized counts over a set of consensus peaks from BigWig tracks per cell type. When using topic modeling, either a regression or classification model can be trained. The regression model uses peak-topic probabilities as target values. The classification model on the other hand first binarizes peak-topic probabilities, and uses the class, or topic, for each peak as target value, as was done in previous work^13,21,29,30^. When using peak heights, a scalar value per cell type is retrieved across each consensus peak (max, mean, sum, or the logarithm of the sum), similar to the scalar value used by ChromBPNet models^8^. These scalar values over the set of cell types are used as targets for multi-class modeling. To make peak heights comparable across cell types, appropriate normalization is necessary. Regular CPM normalization accounts for the different number of cells and sequencing depths across cell types and samples. However, it assumes that each cell type has a similar number of genomic peaks. In other words, using only CPM normalization, a cell type with the same number of cells as another cell type, but with more peaks, will have overall lower values. We account for this issue by rescaling the data based on constitutive peaks. To identify constitutive peaks for scaling, we consider per cell type the highest peaks that are generally accessible (Gini index below the difference between the mean of Gini indices of all peaks minus one standard deviation), under the assumption that these should be of equal height across all cell types, calculate their average height and scale the data such that these constitutive peaks have the same values across cell types (fig. S1a). This peak-scaling normalization is fundamentally a min-max normalization similar to the ReadsInTSS option in ArchR^22,38^, but is applicable directly on BigWigs without the need for the source fragment files.

In both modes, either using topics and peak heights, cell type-specific regions can be selected for training. Such regions can be manually defined by subsetting the consensus peaks, by calculating them through a scATAC-seq data analysis pipeline of choice, or in the case of peak regression they can be calculated from all consensus peaks by selecting regions that have a high Gini index. Here, this is defined as a Gini index greater than the mean of Gini indices of all all regions plus one standard deviation. In the final step, regions are split into a training, validation, and test set, either by splitting on chromosomes or by randomly dividing regions according to user-defined proportions.

For training a classification model, binary cross entropy is used as a loss function. In regression modeling^22,29^, the sum of the average cosine similarity and the log mean squared error (MSE) between the region predictions over all cell types and their target values is set as the default loss function. Optionally, both parts of the loss function can be scaled dynamically during training. Multiple architectures can be chosen, all of which are inspired by previously validated models^6,10,11,42^ (Fig. 1). We suggest to pretrain models on a large set of regions, and fine-tune using cell type-specific regions in a second step. This ensures that the model has seen a wide variety of regions and has an understanding of generally accessible regions but is in the end optimized for regions that uniquely identify topics or cell types that give us insight into their unique enhancer codes. Additionally, the large-scale Enformer or Borzoi foundation models^10,15^ can be used as pre-trained models, whereby CREsted provides transfer learning steps to scATAC data obtained for a specific biological system.

The main purpose of CREsted models is to obtain insights into the underlying model reasoning. CREsted allows for explanations at nucleotide-level resolution, using gradient-based methods^43^ and *in silico* mutagenesis (ISM)^3,6^. For any genomic or synthetic region, a trained model can be used to investigate which motif instances underlie its prediction of accessibility in a query topic or cell type. To perform this task genome-wide, we calculate the enriched patterns for the top cell type-specific regions per model class through tfmodisco-lite^44,45^ and tangermeme^46^. Next, cell types or topics can be clustered based on their best patterns^29^ followed by matching those patterns to TF candidates by using matched scRNA-seq data (if available) and a library of motif-to-TF matches^39^.

As a final module, CREsted provides various methods for the design of synthetic enhancers, as described by Taskiran *et al*.^31^. Sequences can be designed from scratch with *in silico* evolution (ISE), or by the optimal implantation of instances of motifs identified in the previous module. We additionally added a novel optimization function that makes use of the L2 distance, matching the output prediction vector to a predetermined target. This helps to ensure cell type-specificity, by designing cell type-specific sequences while keeping the enhancer off in other cell types.

### CREsted provides detailed insights into enhancer codes of mouse cortical cell types

As a use case for CREsted, we investigated a scATAC-seq dataset^47^ of the mouse cortex that was recently used to benchmark the prediction of *in vivo* enhancer activity^22^. We used cell types at the subclass level (Fig. 2a), obtained pseudobulk chromatin accessibility tracks per cell type and applied the default preprocessing with peak-scaling. We validated our cell type-specific peak scalars by comparing them against the average peak heights from 3,985 promoters from mouse housekeeping genes^48^. As an example, we show the peak height distributions of the promoters of *Canx* and *Zfp106*, two examples that are among the set of the 22 most stable promoters (fig. S1b). This highlights that there is a substantial amount of variability in peak heights across cell types with a coefficient of variation (CV) of 0.48 and 0.37 respectively in CPM normalized peaks. Particularly, peaks from excitatory neurons are lower than those from non-neuronal cell types, with up to a threefold difference in Layer (L) 5 extra-telencephalic (ET) and microglia (Micro-PVM) peak heights. To further generalize these findings, we calculated the average accessibility over all types for all housekeeping gene promoters. While the average expression of all of these genes across cell types is indeed consistent (CV = 0.08) (fig. S1c), their average accessibility is highly variable (CV = 0.38). The variability in peak heights of these housekeeping promoters is indeed consistent with the peak-scaling scalars we identified through CREsted (fig. S1d). In other words, they follow the expected values needed for scaling the housekeeping promoters to the same value.

**Fig. 2:**
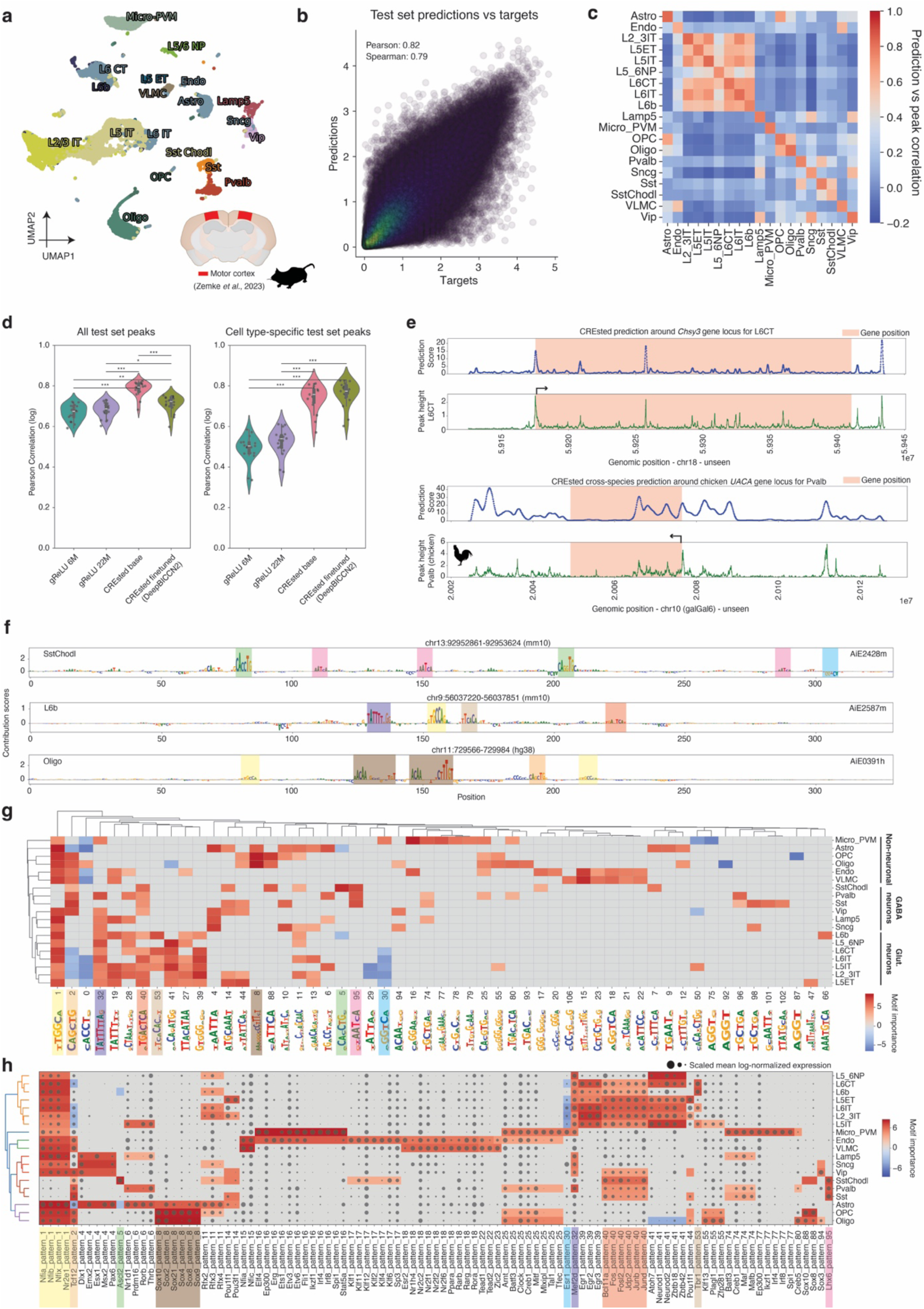
CREsted applied to mouse motor cortex scATAC-seq data. (**a**) scATAC-seq UMAP of the Zemke *et al.*^47^ mouse motor cortex data. (**b**) Scatter plot of log-transformed predicted peak heights and scATAC target peak heights for test set regions over all cell types, generated with *crested.pl.scatter.class_density*. (**c**) Heatmap of Pearson correlations for log-transformed cell type-specific test set peak heights and predictions, separated per cell type, generated with *crested.pl.heatmap.correlations_predictions*. (**d**) Comparison of Pearson correlation between log-transformed peaks and predictions per cell type (n=19) for gReLU and CREsted models, on all test set consensus peaks (left) and cell type-specific peaks (right). Stars indicate adjusted P-values (two-sided t-test, Benjami-Hochberg corrected, * < 0.05, ** < 0.01, *** < 0.001). (**e**) Gene locus scoring with the fine-tuned CREsted model on a gene located in a held-out chromosome for the L6CT class, compared to its scATAC-track (top), and cross-species scoring on a chicken gene locus for the Pvalb class, compared to its chicken scATAC-track^29^ (bottom), generated with *crested.pl.hist.locus_scoring*. (**f**) Contribution scores of three *in vivo* validated enhancers for their corresponding classes, generated with *crested.pl.patterns.contribution_scores*. Colored boxes were added for different identified motifs. Groups of non-neuronal cell types, GABAergic (GABA) neurons and glutamatergic (glut.) neurons are highlighted. (**g**) Clustermap of identified motifs in top 2000 regions per cell type. The color scale indicates motif importance, represented by the log-transformed pattern count. Negatively contributing motifs were given negative importance values. This plot was generated with *crested.pl.patterns.clustermap_with_pwm_logos*. Colored boxes were added for matching motifs found in (f). (**h**) Clustermap of scaled mean log-normalized TF expression over cell types, indicated by dot size. TFs have matching binding sites with a motif in (g). Motif importances for the matched patterns are indicated by color. This plot was generated with *crested.pl.patterns.clustermap_tf_motif*. Colored boxes were added for matching motifs in (f) and (g).

To predict chromatin accessibility levels across cell types, we trained DeepBICCN2, a CREsted peak regression model. We first trained on all 440,993 consensus peaks in the dataset followed by further fine tuning to 73,326 regions with variable chromatin accessibility. On test regions from held-out chromosomes, we obtain an average Spearman correlation over cell types of 0.79, and an average Pearson correlation coefficient (*r*) of 0.82 between log-transformed predictions and peak heights (Fig. 2b). Comparing predicted chromatin accessibility on peaks in the test set to the ground truth shows that predicted cell types always have the highest correlation to their targets (Fig. 2c). Cell types that have similar accessibility profiles, such as glutamatergic neurons (fig. S2), also exhibit high prediction correlation. We compared the predictions of our base- and fine-tuned model against a default (6 million parameter) and large (22 million parameter) model trained with the gReLU framework^35^. Both CREsted predictions on all test peaks and predictions on cell type-specific test peaks significantly outperformed both the base- and the fine-tuned gReLU models (P-value < 0.001) (Fig. 2d).

Encouraged by the strong performance of the DeepBICCN2 CREsted model to predict cell type-specific accessibility of short sequences, we further extended predictions to genomic loci utilizing a sliding window. As an example, we evaluated the *Chsy3* gene, which is specifically expressed in L6 cortico-thalamic (CT) cells. We scored its locus by scoring all regions in that locus for the L6CT class, with a 100 bp step (Fig. 2e). We observed a high correlation (*r* = 0.75) between the predicted chromatin accessibility track and the ground truth, indicating that DeepBICCN2 not only generalizes well to unseen peak regions, but also to inaccessible regions between peaks. We then extended this concept of locus-scoring to genomes of other species. For example, we have recently shown that the enhancer code of mouse and bird interneurons are strongly conserved^29^. To illustrate this with CREsted, we scored the chicken *UACA* gene locus, a gene specifically expressed in Parvalbumin (Pvalb) interneuron cells, with the mouse Pvalb class, and found a strong correlation (*r* = 0.62) between the mouse prediction and chicken scATAC track. This illustrates the possibility of scoring genomic loci in species without scATAC-seq data to identify and decode candidate enhancers.

Next, we used DeepBICCN2 to score 171 *in vivo* validated functional and cell type-specific enhancers^23^. We calculated precision and recall on a set of these enhancers in a multilabel classification setting^22^ and obtained an average precision of 0.77 and recall of 0.93 (fig. S3a and b). We highlight three example enhancers (AiE2428m, AiE2587m and AiE0391h) that have a predicted level of chromatin accessibility that is both strong and specific for the cell type in which these enhancers are also active (fig. S3c and d). The contribution scores of these enhancers highlight a variety of different candidate TFBS (Fig. 2f). To find TF candidates for these TFBS, we calculated contribution scores for the 2000 most specific regions per cell type with *tfmodisco-lite*^44^, clustered frequently occurring patterns based on motif similarity and counted instances; or seqlets, of those pattern-clusters in regions across cell types (Fig. 2g). This results in clustering of cell types based on motif importance and shows logical grouping of non-neuronal, glutamatergic and GABAergic cell types. We then used these globally identified motifs to match them with TF candidates using the SCENIC+ motif-to-TF database^39^. To further filter the list of candidates, we used corresponding scRNA-seq data^47^ and filtered to TFs that followed a similar expression profile compared to the motif importance across cell types. We find previously observed^29^ specific factors for deep layer glutamatergic neurons such as TBR1 and NFI, RFX3 and EGR for upper layer glutamatergic neurons, SOX10 and CREB5 for oligodendrocytes (Oligo), SPI1, IRF and MAFB for microglia/perivascular macrophages (Micro-PVM) and LHX6 and MAFB for medial ganglionic eminence (MGE) interneurons. A set of TFBS for these TFs are also found back in the validated enhancers described above (Fig. 2f). Interestingly, we identified an E-box motif with the consensus sequence CAGGTG that is unique to somatostatin-chondrolection (SstChodl) cells, with ASCL2 as a potential TF candidate. Mutating such motif instances in the SstChodl AiE2428m enhancer to the more canonically found CAGCTG motif in all interneurons, changes its CREsted predictions to be generally accessible in MGE interneurons (fig. S3e and f). This highlights the high resolution of CREsted models, from finding global motifs representing cell type-specific enhancer code, to capturing single nucleotide variations in TFBS that are strongly predicted to affect cell type-specific accessibility.

Overall, this example represents a complete run through the CREsted pipeline, from data preprocessing to enhancer code analysis. It highlights that CREsted models are capable of interpreting the enhancer code of mouse motor cortex cell types at high resolution, with strong predictive capabilities at the region and gene locus level and in depth capturing of the motifs that underlie the predicted cell type-specific chromatin accessibility.

### A CREsted human PBMC model captures validated TFBS

To illustrate the capability of CREsted models to also function in other tissues and species, we trained DeepPBMC, a peak regression model on human PBMC data^49^. Following the same training procedure of the mouse cortex model, we pre-trained our model on a set of 278,687 consensus peaks, followed by fine-tuning on 51,644 cell type-specific peaks. On a set of held out specific peak regions, the model obtains an *r* of 0.71 (fig. S4).

First, we used CREsted to generate a DNA-sequence embedding from DeepPBMC of the 1,000 most specific peaks per cell type, based on the combined highest differential peak and prediction score (Fig. 3a, fig. S5). This demonstrates that the model can internally represent enhancer codes of PBMC types in a distinguishable manner. Next, we assessed the ability of CREsted to identify validated TFBS. For this purpose, we investigated an enhancer for the *CD79A* gene, expressed in B-cells, and an enhancer for the *TCRα* gene, which is expressed in T-cells. For both enhancers, TFBS were previously identified and experimentally validated^50,51^. (Fig. 3b). Indeed, CREsted finds the validated TFBS for PAX5, SP1, EBF1, RUNX1 and E2A in the *CD79A* enhancer and SP1, ATF/CREB, TCF/LEF, CBF and ETS in the *TCRα* core enhancer. Furthermore, CREsted also proposes additional TFBS, such as for IRF8 and REL in the *CD79A* enhancer and an additional ETS-like TFBS upstream of the *TCRα* core enhancer. These additional TFBS were also found for relevant classes of the Borzoi model (fig. S6A and B). Next, we investigated whether DeepPBMC is also capable of capturing the TFBS of highly complex enhancers, such as the classical Interferon-β (*IFNB1*) enhanceosome that is active in dendritic cells. This enhancer consists of multiple overlapping TFBS in a 50 bp window and has been structurally resolved together with all bound TFs^52,53^. By investigating contribution scores on both strands of this enhancer, we find that the model retrieves a large part of the enhanceosome’s complexity, only missing TFBS for p50, c-Jun and one of the four IRF TFBS (Fig. 3c). Compared to two classes in Borzoi that represent dendritic cells, the DeepPBMC identifies substantially more (34%) important nucleotides within this enhanceosome (fig. S6C). These findings suggest that DeepPBMC identifies validated TFBS in known enhancers, even in complex regulatory systems such as enhanceosomes.

**Fig. 3:**
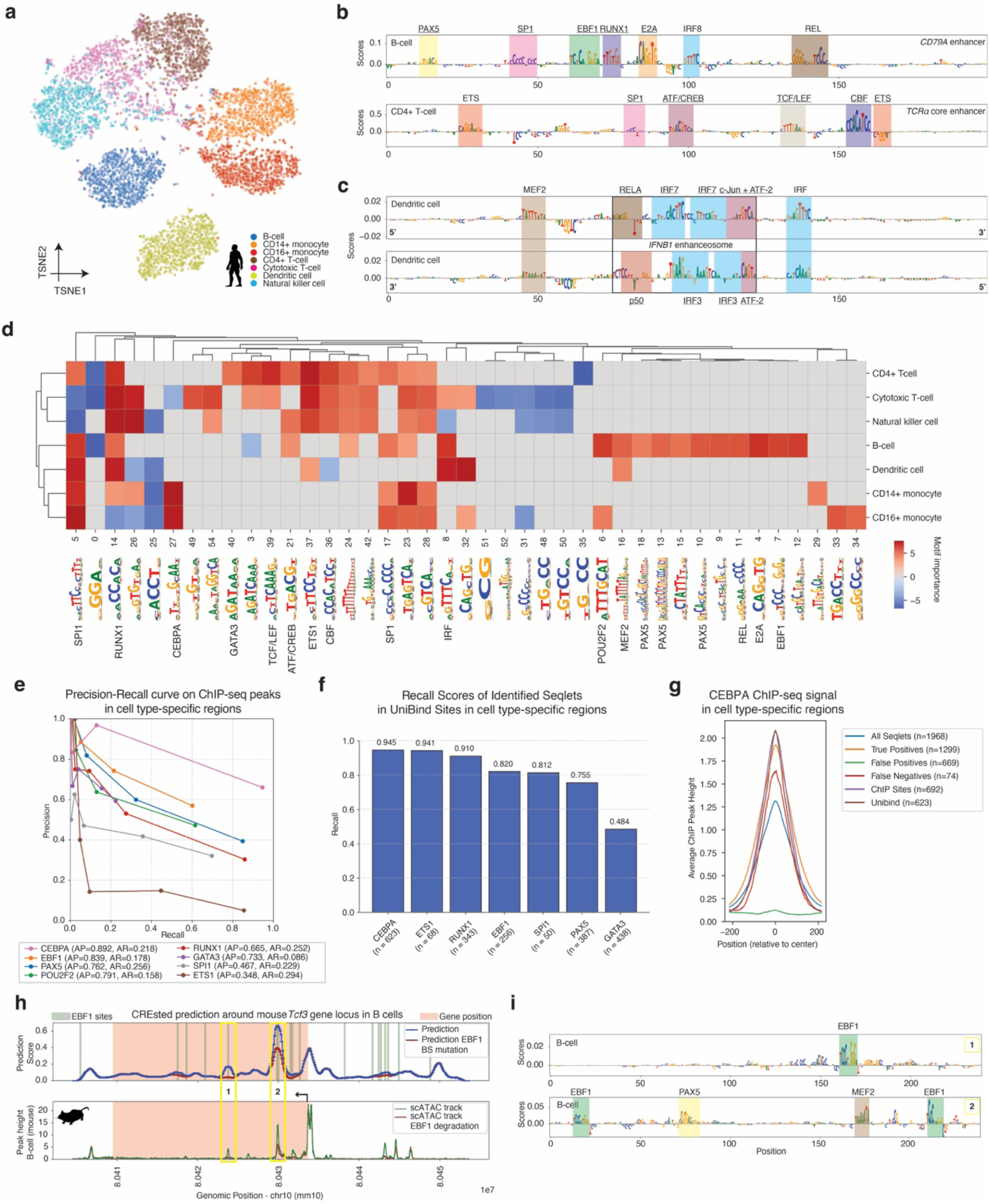
A human PBMC CREsted model identifies functional TFBS. (**a**) tSNE of region model embeddings for the top 1,000 specific regions per PBMC type. The assigned class determines the coloring. (**b**) Contribution scores of the *CD79A* (hg38 chr19:41,876,056-41,878,170) (top) and *TCRα* core (hg38 chr14:22,555,403-22,557,517) (bottom) enhancers for the B-cell and CD4+ T-cell class respectively. The center 200 bp of both enhancers is shown. (**c**) Contribution scores for both strands of the *IFNB1* enhanceosome (hg38 chr9:21,076,963-21,079,077, zoomed to center 200 bp) for the dendritic cell class. The enhanceosome’s location is highlighted with the black box. In (b) and (c), colored boxes indicate found motifs. Validated TFBS are underlined, proposed TFBS are not. (**d**) Clustermap of identified motifs in top 1,000 regions per cell type from (a). The color scale indicates motif importance, represented by the log-transformed pattern count. Negatively contributing motifs were given negative importance values. Motifs were manually annotated with the results from (b) and (c) and motif databases. (**e**) Precision-recall curve on comparing model-identified seqlets and ChIP-seq peaks in the top 1,000 peaks for the corresponding cell type. Thresholding is done on the average seqlet contribution score. Average precision (AP) and recall (AR) over the thresholds are indicated in the legend. (**f**) Bar plot of recall of identified seqlets inside UniBind sites in the top 1,000 peaks for the corresponding cell type. (**g**) Average ChIP peak height of different sets of proposed TFBS for CEBPA in the top 1,000 most-specific CD14+ monocytes. (**h**) Cross-species scoring on the mouse *Tcf3* gene locus in B-cells before (blue) and after (red) EBF1 TFBS mutation (top), compared to mouse scATAC tracks before (green) and after (red) EBF1 degradation in mouse precursor B-cells^63^. (**i**) Contribution scores for two highlighted regions before binding site mutation in (h) for the B-cell class. EBF1 sites are highlighted in green, other proposed PAX5 and MEF2 TFBS are also highlighted.

To obtain a more global view of cell type-specific enhancer codes in PBMC types, we ran CREsted’s pattern clustering analysis and identified *de novo* motifs for core sequence patterns that determine cell type-specific region accessibility in the top 1,000 regions per cell type (Fig. 3d). All of the TFBS identified in Fig. 3b and c are found back in their corresponding cell types. Motifs are identified for previously established key TFs, such as EBF1, PAX5 and POU2F2 in B-cells^39^ and ETS1, RUNX1 and GATA3 in T-cells^54–56^ and CEBPA and SPI1 in monocytes^57,58^.

Next, we asked to what extent the instances of the identified seqlets overlap with actual functional TFBS. To answer this, we used publicly available chromatin immunoprecipitation sequencing (ChIP-seq) data for a selection of key TFs (EBF1, PAX5, POU2F2 in B-cells, CEBPA and SPI1 in CD14+ monocytes, and RUNX1, ETS1 and GATA3 in CD4+ T-cells^56–60^), and evaluated the overlap of identified seqlets with ChIP-seq peaks for their corresponding TFs. Overall, we observe a high precision (average precision over all thresholds for all TFs of 0.75) for nearly all TFs and this precision increases with prediction score (Fig. 3e). The recall is lower overall, which can be attributed to several factors, such as false positive (phantom) ChIP-seq peaks, ChIP-seq peaks with indirect TF binding^61^, the difference between the cell line used in the ChIP-seq assay and the PBMC data, and the parameters used for tfmodisco-lite that can influence seqlet-calling drastically. To avoid these pitfalls, we resorted to UniBind^62^ sites, which represent directly bound ChIP-seq peaks, and examined if at UniBind locations a seqlet was observed. This resulted in a much stronger recall, with an exception for GATA3, both inside specific peaks and in all consensus peaks (Fig. 3f, fig. S7). The high concordance between predicted TF binding site and ChIP-seq signal can also be observed by aggregating ChIP-seq signal over all seqlets (Fig. 3g for CEBPA, fig. S8 for other TFs), and that false positive (FP) seqlets without ChIP-seq peak still have low ChIP-seq signal, suggesting the presence of weak TFBS (fig. S9). These analyses indicate that seqlets identified by CREsted models strongly overlap with ChIP-seq peaks and UniBind sites for a variety of cell types, with an increase in confidence related to prediction strength.

This functional TF binding onto CREsted-predicted TFBS is further confirmed by simulating the effect of TF degradation. Particularly, we scored the mouse *Tcf3* locus for the B-cell class and then rescored that locus with all instances of EBF1 TFBS mutated to become dysfunctional. We then compared the predicted chromatin accessibility profiles to the profile in mouse precursor cells in which the EBF1 protein was acutely degraded^63^. We observe an *r* of 0.55 and 0.60 respectively for control and treated B-cells (Fig. 3h). The two regions in this locus that are most strongly influenced by EBF1 protein degradation indeed contain EBF1 binding sites according to DeepPBMC (Fig. 3i).

These results strongly support that TFBS identified by CREsted overlap with functional sites, both for known enhancer sequences, and globally through the identification of seqlets of essential motifs per cell type.

### CREsted identifies high similarity between mesenchymal-like enhancer codes in cancer

Cancer cell states have been enigmatic to robustly compare between patient biopsies, due to strong patient-specific epigenomes and transcriptomes^64,65^. The same cancer cell states can recur across different tumor types, such as epithelial-mesenchymal-transition (EMT) related or mesenchymal-like cell states reported in melanoma, glioblastoma (GBM), and other cancer types^66–68^. Previously, we modeled enhancers of the mesenchymal-like (MES-like) cell state in melanoma using DeepMEL^13^ and DeepMEL2^69^, with both models now ported to CREsted and available from the CREsted model repository. However, MES-like regulatory programs have to our knowledge not yet been compared between cancer types at the level of enhancer logic. We hypothesized that CREsted models could make abstraction of patient-specific genomic aberrations and enable a direct comparison of regulatory programs active across cancer types. For this purpose, we aimed to compare the MES-like state of melanoma and GBM. Furthermore, cancer is often modeled using cancer cell lines (e.g., for drug target discovery), but cell lines do not necessarily reflect *in vivo* cancer states. To address this, we will compare the enhancer code of GBM cell lines and patient biopsies. This strategy resembles the comparison of regulatory programs between species, for cell types with diverged transcriptomes^29^.

We constructed an ATAC-seq dataset of cancer cell lines, using publicly available tracks for two ENCODE^59,70^ deeply-profiled-cell lines (DPCLs), namely HepG2 and GM12878; three previously generated^39^ melanoma cell lines (two MES-like, MM029 and MM099, and one melanocytic-like, MM001); and newly generated ATAC-seq data for three GBM cell lines (A172, M059J, and LN229). A172 and M059J were previously reported to have a MES-like state, while LN229 was mostly characterized as proneural^71,72^. We trained a peak regression CREsted model on these cell lines, named DeepCCL, using 285,790 regions, and fine-tuned it on 140,761 regions (Fig. 4a). The base DeepCCL model groups together the MES-like states derived from GBM and melanoma, compared to the included - non-MES-like - control group of DPCLs, which is also recapitulated by the actual peak accessibility (fig. S10a). Additional finetuning of DeepCCL can partially distinguish the GBM and melanoma MES-like cell lines but still identifies commonalities in their regulatory programs (Fig. 4b, fig. S10b). To confirm our model findings, we added an ensemble of ChromBPNet models trained separately on the same cell lines. The ChromBPNet ensemble illustrated a similar grouping of the MES-like states across the GBM and melanoma cell lines (fig. S10a and b), despite its lower performance on test data compared to the DeepCCL model (fig. 4c). We also examined the previously validated IRF4 melanocyte-like enhancer, for which the CREsted-based ISM achieved a higher Spearman correlation (0.75) with in vitro mutagenesis values^73^, compared to ChromBPNet (0.67) and DeepMEL2 (0.66) models (fig. S10c). To further investigate the similarities between the MES-like states, we compared nucleotide contribution scores in a candidate *AXL* enhancer^69^ (Fig. 4d) that is accessible in both GBM and melanoma MES-like lines and is explained in both by a combination of AP-1, TEAD and ZEB motifs. Contribution scores from the ensemble ChromBPNet model and DeepMEL2 further confirm these motifs (fig. S10d). Motif analysis confirms similar cis-regulatory logic between these cell lines, despite their different tissue of origin, with main contributions of AP-1, TEAD, RUNX, NFI, and ATF/CREB motifs (Fig. 4e). The fine-tuned model reveals a bias towards TFAP2 and MITF motifs for the melanoma MES-like state, suggesting that a remnant melanocytic-derived program is still active; this was previously difficult to extract when comparing with melanocyte-like melanoma lines alone^74,75^.

**Fig. 4:**
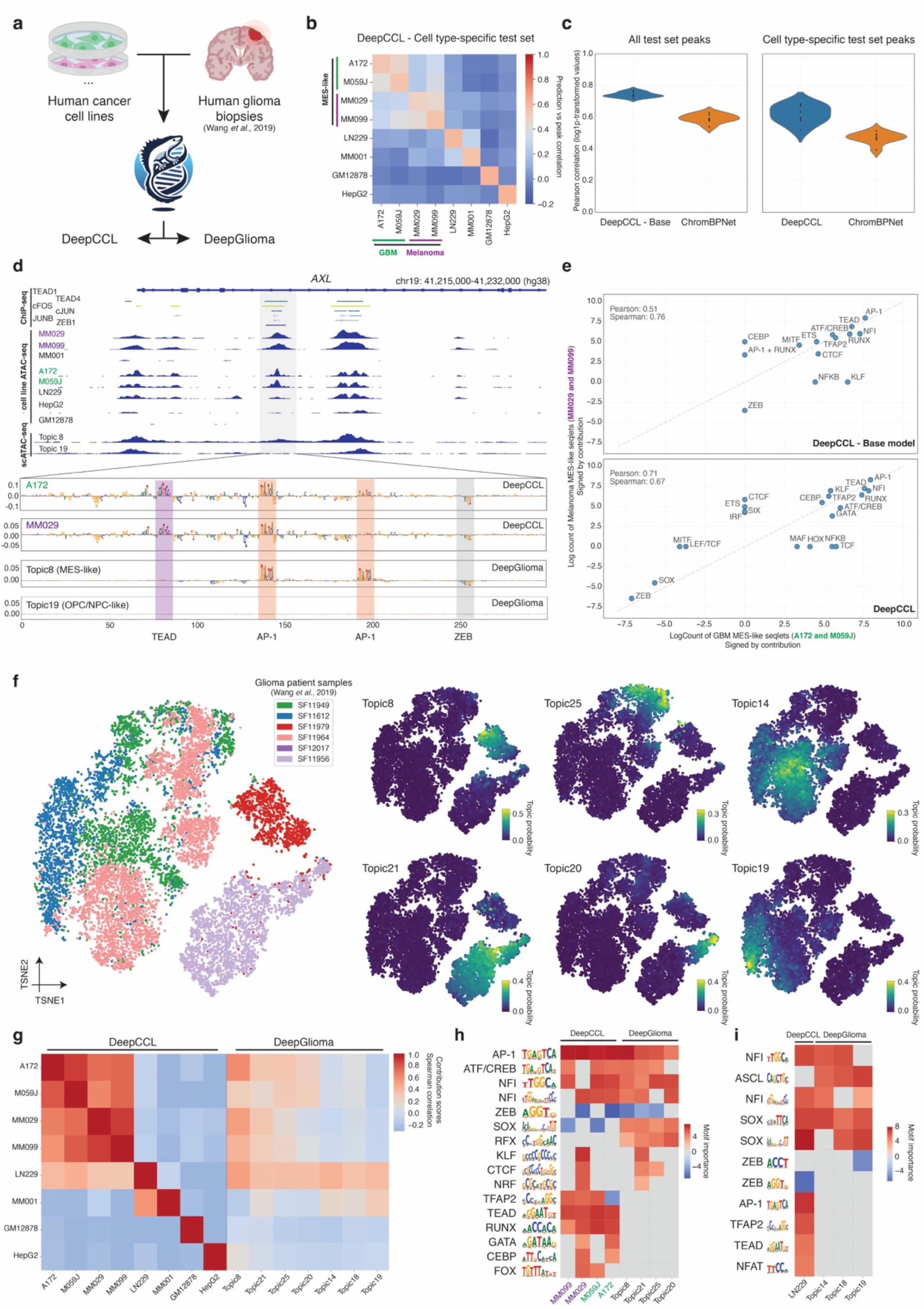
Comparing MES-like states across cancers in cell lines and biopsies. (**a**) Overview of the DeepCCL and DeepGlioma model trained on human cancer cell lines and GBM biopsy data respectively. (**b**) Heatmap of Pearson correlations for log-transformed cell type-specific test set peak heights and predictions from DeepCCL, separated per cell line. MES-like states are highlighted. (**c**) Comparison of Pearson correlation between log-transformed peaks and predictions per cell line (n=8) for DeepCLL (base model and fine-tuned) and an assembled ChromBPNet model, on all test set consensus peaks (left) and cell type-specific peaks (right) (**d**) Highlighted example region intronic in *AXL*, showing multiple ChIP-seq, cell line ATAC tracks and biopsy topic scATAC tracks (top). Contribution scores for MES-like model classes and an OPC/NPC-like class from the DeepCCL and DeepGlioma models, with ChIP-seq matched motifs highlighted. (**e**) Comparison of log counts of CREsted-identified grouped motif sets between GBM MES-like and melanome MES-like cell lines for the DeepCCL base model (top) and DeepCCL (bottom). Negatively contributing patterns were given a negative count. (**f**) tSNEs based on pycisTopic’s topic-cell contributions of human biopsy cells, colored by patient sample (left) and topic probability for a set of topics (right). (**g**) Contribution score Spearman correlation between contribution scores obtained from the DeepCCL and DeepGlioma model per model class for the top 1,000 most specific regions per cell line and per topic. (**h**, **i**) Heatmap of identified motifs in top 2,000 regions for DeepCCL and DeepGlioma classes. The color scale indicates motif importance, represented by the log-transformed pattern count. Negatively contributing motifs were given negative importance values. Motifs were manually annotated.

Next, to investigate whether this MES-like program observed in cell lines is also active in tumor biopsies, we re-used a previously published scATAC-seq dataset of human gliomas^76^. To avoid any biases in cell type annotation, we used pycisTopic^41^ to identify topics across this scATAC-seq atlas, yielding both patient-specific topics and shared topics across patients (Fig. 4f, fig. S11a). Next, we trained a CREsted topic classification model on these data, named DeepGlioma, that predicts chromatin accessibility across GBM biopsy states (Fig. 4a, fig. S11b). By comparing cell line accessibility and predictions from DeepCCL with topic accessibility, we identified a set of potential MES-like biopsy topics and three candidate OPC/NPC-like topics (fig. S11c). To enhance this comparison^29^ and enable its abstraction away from non-sequence-mediated changes, such as copy number variations, we used DeepGlioma to compare nucleotide contributions with those obtained by DeepCCL focusing on the MES-like topics (Fig. 4g). This revealed one biopsy topic (Topic 8) with a high similarity to the MES-like cell line enhancer code, but clearly lower compared to any of the pairwise similarities found between the cell lines. In other words, the similarity between MES-like cell lines derived from GBM or melanoma is higher than the similarity between GBM cell lines and GBM biopsies. Clustering patterns found by both models revealed AP-1 and CREB/ATF motifs to be shared between the MES-like cell lines and the MES-like biopsy topic, while TEAD motifs are specific to cell lines, and SOX and RFX motifs are specific to biopsies (Fig. 4h). The cell line specific importance of TEAD is further illustrated by the AXL enhancer in Fig. 4d. On the other hand, the previously characterized proneural cell line LN229^71^ shows similarity to OPC/NPC-like cancer cells in GBM tumors (Fig. 4g), sharing the SOX monomer/dimer motifs (Fig. 4i). Also here, certain motifs are missing from the cell line enhancer logic, such as ASCL. Notably, LN229 still shares a part of the identified MES-like program (e.g., AP-1, TEAD) making it more accurately defined as a mixed proneural-mesenchymal cell line from the chromatin accessibility perspective^75^. In conclusion, this case study illustrates how CREsted models can be used to compare cancer cell states across biopsy samples and cell lines, highlighting that the often-found MES-like enhancer logic in cell lines is not fully recapitulated in the tumor.

### CREsted-trained models outperform large, pre-trained models on cell type-specific chromatin accessibility predictions

Besides training enhancer models on specific scATAC-seq data from scratch, an alternative strategy involves transfer learning from large pre-trained models, such as Enformer^10^ and Borzoi^15^. In this strategy, there may be a benefit from the large knowledge base these models are trained on, which could be transferred to the dataset-specific models^25,26,77,78^. The CREsted package provides the option to evaluate these models as they are, and to use these pre-trained models for transfer learning. For this latter purpose, CREsted provides modules to update their architecture, to accommodate sequences of any specified length as input, and to add an extra dense layer after the final embedding to predict peak heights for the set of cell types in the new scATAC-seq data (Fig. 5a).

**Fig. 5:**
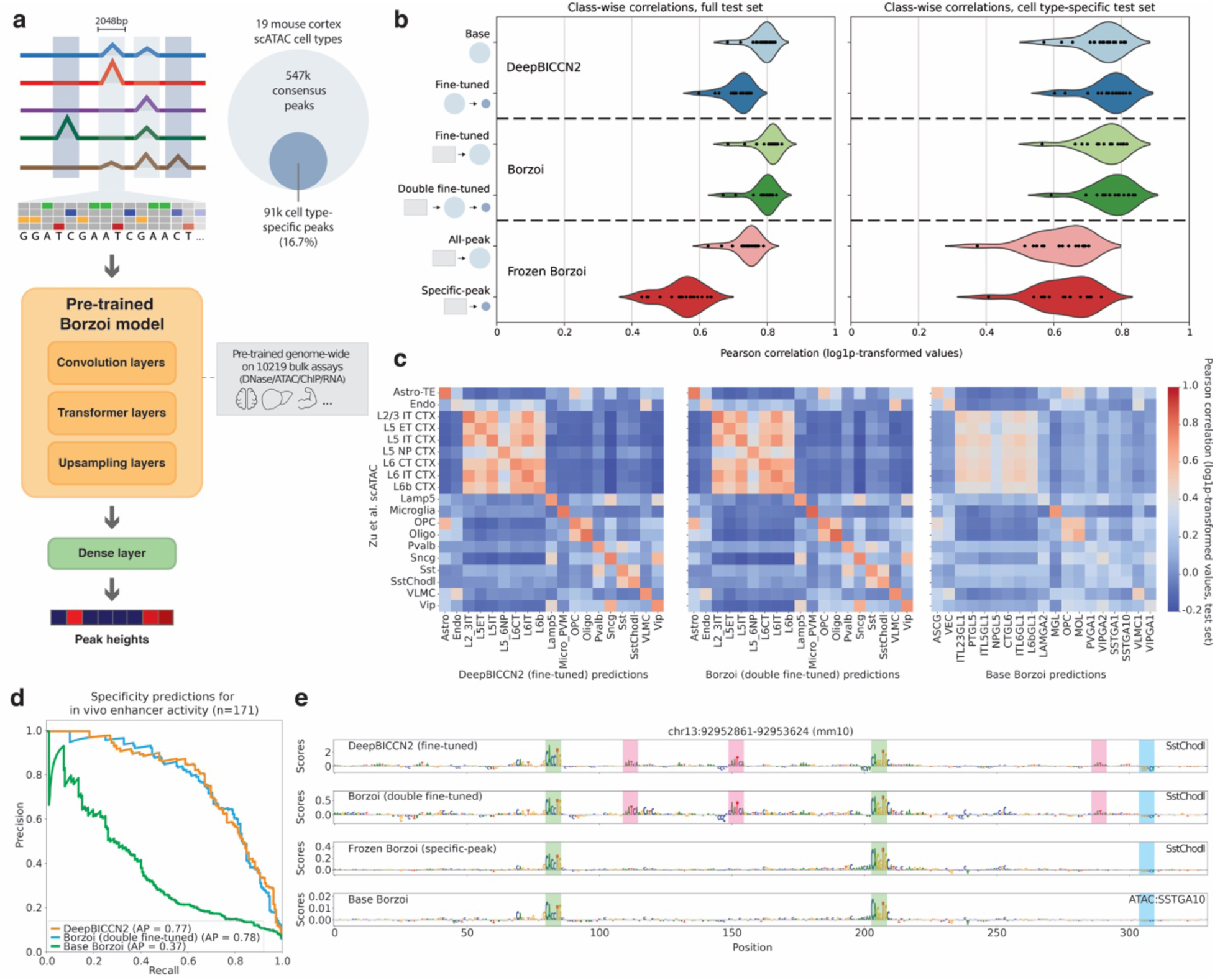
Transfer learning from and benchmarking against a large pre-trained sequence-to-function model. (**a**) Schematic overview of the Borzoi transfer learning process, fine-tuning and evaluating a modified Borzoi model on all consensus peaks, cell type-specific peaks, or both. (**b**) Comparison of mean *r* between log-transformed predictions and peak heights per cell type (n=19) with the test dataset for the DeepBICCN2 model (Fig. 2), the fine-tuned Borzoi, and a fine-tuned Borzoi with the model trunk (‘Pre-trained Borzoi model’) frozen. Icons denote the datasets these models were successively trained on (original Borzoi training data, mouse cortex consensus peaks, or mouse cortex cell type-specific peaks), matching (a). (**c**) Heatmaps showing the correlation of model predictions with peak heights for matching classes from the Zu *et al*. mouse brain atlas^16^, evaluated on the cell type-specific peaks in each model’s corresponding test set. **(d)** Precision-recall curve on specificity predictions on a set of *in vivo* validated enhancers. **(e)** Contribution scores for a validated SstChodl enhancer^23^ (AiE2428m) for the specific models and a matching class from the base Borzoi model.

To illustrate this training method, we used mouse motor cortex scATAC-seq data^47^ also used to train DeepBICCN2 (Fig. 2) and employed the Borzoi model as a starting model from which we transfer learned to the motor cortex cell types using 2,048 bp input sequences. Like in the recommended CREsted approach for training models from scratch, the model was first fine-tuned on the entire consensus peak training set (440,993 regions) and further fine-tuned on a training set of cell type-specific peaks (73,326 regions) (Fig. 5a). Fine-tuning solely on the cell type-specific peaks showed worse performance compared to the double fine-tuning approach (Fig. S12a). As a reference model, we also trained a version of this model with the pre-trained section of the model frozen, training only the new dense layer on either the consensus regions or only the specific set.

Next, we compared DeepBICCN2 with the different Borzoi-based fine-tuned models by evaluating all models on the held-out test splits of the full consensus peak and the cell type-specific peak datasets (Figs. 5b, fig. S13). When evaluating the models on the consensus peak test set, the base DeepBICCN2 and fine-tuned Borzoi model achieved similar levels of performance, 0.79 and 0.80 mean class-wise *r* respectively, with the all-peak frozen Borzoi model performing slightly worse (0.73 mean *r*). The models further fine-tuned on the cell-type specific training set all decreased in performance on the consensus peak set. However, fine-tuned Borzoi’s performance on the full test set only dropped very slightly after double fine-tuning (0.80 to 0.78 mean *r*), while DeepBICCN2’s performance drop-off is more substantial (0.79 to 0.71 mean *r*), potentially indicating a better capability of large pre-trained models like Borzoi to retain predictive capabilities for non cell type-specific peaks.

On the cell type-specific test set, the DeepBICCN2 and fine-tuned Borzoi models each realize a slight performance boost after the second round of fine-tuning on the cell type-specific training set (0.74 to 0.76 mean *r* and 0.75 to 0.78 mean *r* respectively), effectively achieving very similar performance on this set overall (Fig. 5b). The frozen Borzoi models clearly perform worse than either fully trained model type, reaching 0.61 mean *r* for the all-peak model and 0.63 mean *r* for the specific-peak model. We conclude that fine-tuning on Borzoi models requires full re-training, rather than training a single fully connected layer on the Borzoi embeddings; and that simple CREsted architectures with only 6M parameters and trained on 19 chromatin accessibility tracks match the performance of large models like Borzoi with 170M parameters, trained on 6230 regulatory and 3989 gene expression tracks.

To further validate the performance of the new Borzoi and DeepBICCN2 models as compared to the base Borzoi model, we evaluated the model’s performance against cell types from an external dataset. This dataset consists of cortical cell types from a whole mouse brain chromatin accessibility atlas, which match with the cell types of the mouse motor cortex^16^ (Fig. 5c). Borzoi was trained to predict chromatin accessibility tracks of similar cell types, but from another mouse brain atlas^79^. The 159 tracks from this earlier atlas, together with 314 other scATAC pseudobulk classes, form a minor part of the regulatory tracks used for base Borzoi pre-training (4.6% of all tracks). Comparing these matching classes shows that the fine-tuned Borzoi and DeepBICCN2 have similar performance and outperform the base Borzoi model when generalizing to new measurements of the same biological cell types (Fig. 5c). While the base Borzoi model scores lower on all classes, it particularly has difficulty predicting the different subclasses of GABAergic and glutamatergic neurons. Although the glutamatergic neurons in particular have similar chromatin accessibility profiles, correlations between the measured peaks show these subclasses are still distinct (Fig. S14).

Furthermore, we calculated precision and recall on the set of 171 *in vivo* validated mouse enhancers^22,23^ (Figs. 4d, fig. S15) as was done in fig. S3B. Similar to Fig. 5b and c, DeepBICCN2 and the double fine-tuned Borzoi model are at the same level with an average precision (AP) of 0.77 and 0.78 respectively, while the base Borzoi model performs substantially worse (AP = 0.37). These results indicate that the base Borzoi model does not perform well at increased cell type resolution and has difficulty differentiating between them, even if those were present during training, and that fine-tuning is necessary to achieve comparable results to models trained from scratch.

Focusing on how fine-tuning affects explainability of the models, we compared the contribution scores of the three models further trained on the specific peaks, as well as the base Borzoi model, on a validated enhancer in SstChodl cells (Fig. 5e), as well as other validated enhancers in other cortex cell types (fig. S16)^23^. We observe that the most important motifs are captured by all models, but that the fine-tuned DeepBICCN2 and the double fine-tuned Borzoi models find additional LHX6-like (Fig. 2h) motifs, which are not found by base Borzoi or the frozen fine-tuned model.

In summary, CREsted allows for training of new models that can outperform large pre-trained models, whether through transfer learning or from scratch. We show that the base large, pre-trained models appear to fail at differentiating cell types at high resolution in the mouse motor cortex.

### CREsted designs enhancers specifically active in targeted cell types of a developing zebrafish

To assess the use of CREsted for the design of organism-level cell type-specific enhancers, we trained a peak regression model on a scATAC-seq dataset of the developing zebrafish embryo^80^ (DeepZebrafish; Fig. 6a). This dataset comprises 20 developmental stages, and 639 cell type-timepoint combinations that were used as separate classes for the model. The model was trained on a set of 793,273 consensus peaks and fine-tuned on 89,637 cell type-timepoint-specific peaks (Fig. 6a). Next, we evaluated the performance of the model to predict the cell type-specificity of 54 validated enhancers^80^, and we observed that indeed 76% of cell type-specific enhancers had a high and specific prediction score for the corresponding cell type-class in the model (Fig. 6b).

**Fig. 6:**
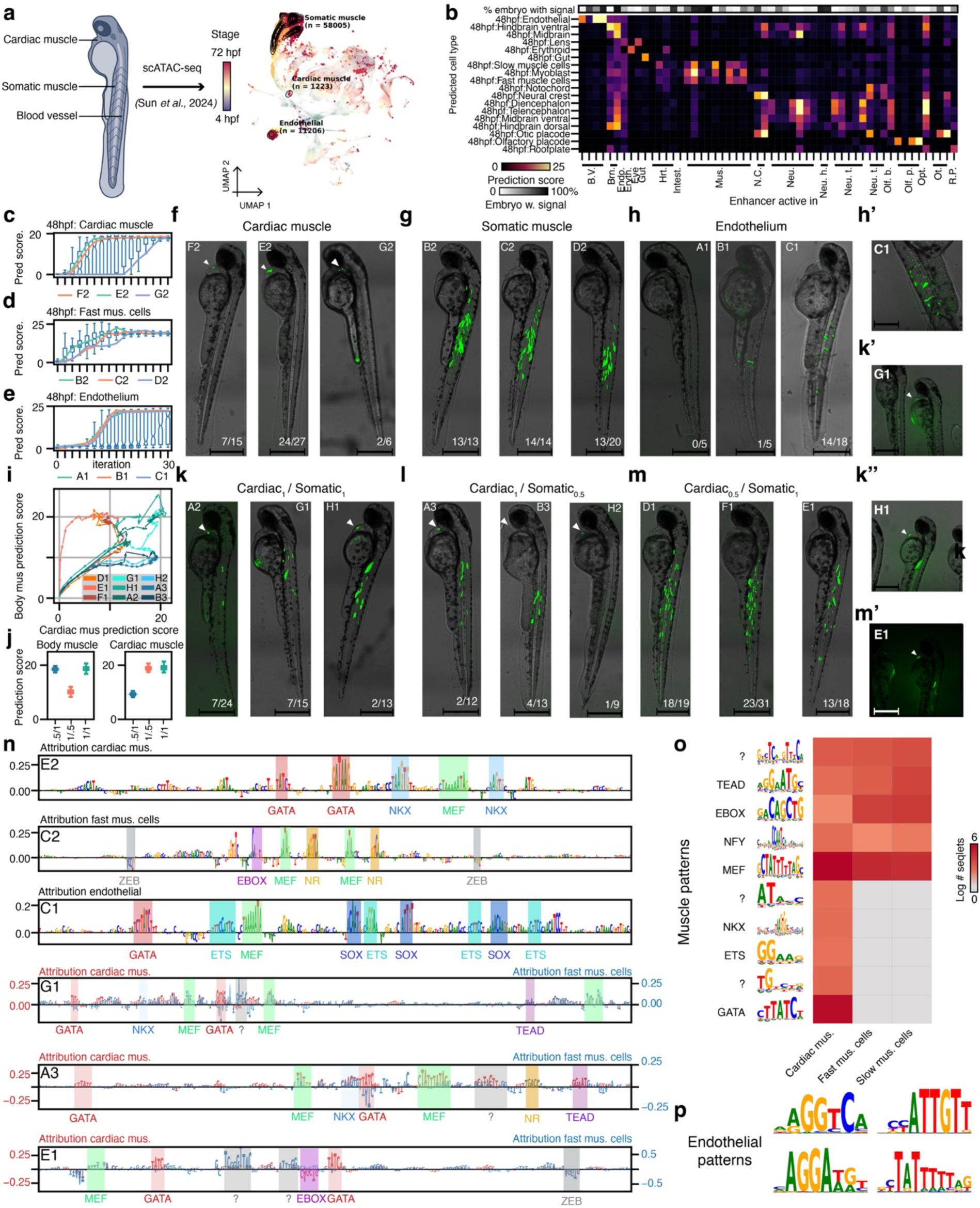
DeepZebrafish can be used to design cell type-specific enhancers in the whole zebrafish over development. (**a**) Schematic overview of 48 hours post fertilization (48 hpf) zebrafish embryo and UMAP of scATAC-seq cells with cell types of interest highlighted (**b**) Heatmap of prediction score of DeepZebrafish on validated zebrafish enhancers and percentage of embryos that were active for the enhancer. (**c-e**) Prediction score on designed enhancers for each iteration of the design process (boxplot; n=200) and prediction score of tested enhancers for each iteration (lines) for enhancer designed for: cardiac muscle (c), somatic muscle (d) and endothelium (e). (**f-m**) brightfield and fluorescent images (overlayed) of enhancer reporter assays in 48 hpf zebrafish embryos for enhancer designed for: cardiac muscle (f), somatic muscle (g) endothelial cells (h), cardiac and somatic muscle in 1 to 1 ratio (k), cardiac and somatic muscle in 2 to 1 ratio (l) and cardiac and somatic muscle in 1 to 2 ratio (m). Scale bar is 500µm. Fractions indicate the number of embryos that look similar to the representative one over the total number of embryos that were imaged per condition. (**n**) contribution scores for one synthetic enhancer per set of enhancers. (**o**) Heatmap of identified patterns in enhancers designed for cardiac and/or somatic muscle cells showing the natural log of the number of seqlets per class over all designed enhancers. (**p**) Patterns identified in enhancers designed for endothelial cells. B.V.: blood vessel; Brn.: Brain; Endo: endothelial; Hrt.: heart; Intest.: intestine; Mus.: muscle; N.C.: neural crest; Neu.: neuron; Neu. h.: neuron(head); Neu. t.: neuron(trunk); Olf. b.: olfactory bulb; Olf. p.: olfactory placode; Ot.: otic; R.P.: roof plate. For box-plots in c-e, and j the top/lower hinge represents the upper/lower quartile and whiskers extend from the hinge to the largest/smallest value no further than 1.5 × interquartile range from the hinge, respectively. The median is used as the center.

Next, we designed enhancers specific for endothelial cells and two closely related cell types: cardiac muscle and somatic muscle cells (the latter represented by the “Fast muscle cells” and “Slow muscle cells” classes in the model; Fig. 6c-e). We used a greedy search, ISE, to optimize randomly initialized DNA sequences (see Methods) towards cell type-specific enhancers exploiting the model as a guiding oracle^31^. To ensure cell type specificity, we defined a cost-function to minimize the Euclidean distance (L2 distance) between the model’s prediction-vector and a hypothetical vector with targeted levels of chromatin accessibility (eq. 1). In this case, we aimed for high levels in the target cell type and no accessibility in other cell types. We set the target predicted level of chromatin accessibility to the average level of predicted chromatin accessibility of the positive validated enhancers.

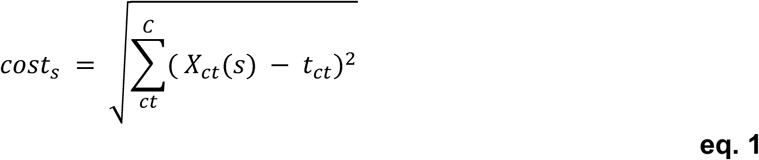

With:

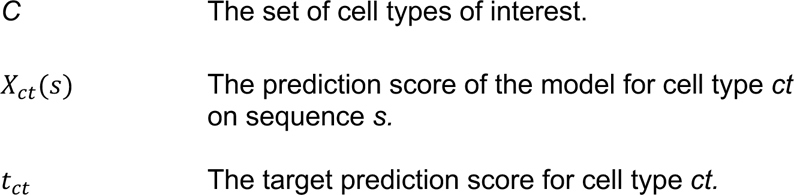

Using this approach, after 30 iterations, most designed sequences reached the target level of predicted chromatin accessibility (Fig. 6c-e). For each target cell type, we tested a set of three candidate enhancers using an *in vivo* enhancer reporter assay in zebrafish embryos. We designed constructs containing the designed enhancer upstream of the minimal ZSP promoter^81^ and an eGFP, with a downstream fluorescent transgenesis reporter (lens-directed cryaa:mCherry). We injected these plasmids with transposase RNA in 1 cell stage oocytes, using Tol2 transposon to integrate the construct in the embryos’ genome and observed enhancer activity at 48 hours post fertilization (hpf). All enhancers designed for cardiac muscle cells were specifically active in the heart (fig. 6f) and all enhancers designed for somatic muscle cells were specifically active in the target cell type (Fig. 6g). The efficiency of cardiac muscle enhancers was overall lower (fig. 6f). Only one of three enhancers designed for endothelial cells had strong and specific activity, another endothelial enhancer was active in the correct cell type, but the signal was weak (Fig. 6h).

Encouraged by this result, we asked if we could use the model to design enhancers to be active in both cardiac and somatic muscle cells and if the level of prediction score has a functional relationship to enhancer activity. To this end, we designed sets of enhancers with equal levels of predicted accessibility in both cell types (cardiac_1_/somatic_1_) and sets of enhancers that had half the level of predicted accessibility in either cardiac (cardiac_0.5_/somatic_1_) or body muscle cells (cardiac_1_/somatic_0.5_). Again, after 30 iterations, most sequences reached their target level (Fig. 6i-j). For all cardiac_1_/somatic_1_ enhancers (Fig. 6k) and most cardiac_1_/somatic_0.5_ enhancers (Fig. 6l) we detected activity in both cardiac and somatic muscle cells, surprisingly the efficiency of these enhancers was overall substantially lower. Both in the sense that less embryos were positive per enhancer and for each embryo that had activity in somatic muscle cells, a lower amount of muscle cells showed reporter expression compared to enhancer designed for somatic or cardiac muscle alone (Fig. 6k-l). Only one out of three cardiac_0.5_/somatic_1_ enhancers was active in both cell types and the two enhancers that were only active in the somatic muscle cells showed overall higher efficiency (with both more embryos being positive and a larger number of somatic muscle cells showing reporter expression; Fig. 6m). From this we conclude that DeepZebrafish can be used to design dual code enhancers but that controlling the magnitude of reporter expression requires further investigation.

Finally, we made use of the CREsted pattern clustering analysis to identify the main sequence characteristics underlying the designed enhancer sets. Both cardiac and somatic muscle enhancers have a set of shared features corresponding to MEF, TEAD and EBOX motifs (Fig. 6n-o). Cardiac muscle enhancers have additional features corresponding to GATA and NKX motifs (Fig. 6n-o). Surprisingly, we could not identify individual motifs that were specific to the somatic muscle enhancers (Fig. 6o). Lastly, for endothelial enhancers we found nuclear receptor, SOX, ETS and MEF motifs to have a positive contribution (Fig. 6n, o).

In conclusion, CREsted scales well to large-scale developmental scATAC-seq datasets and can be used to design enhancers that are specific to a target cell type across an entire organism.

## Discussion

High throughput identification and functional characterization of *cis*-regulatory elements has been a long-standing challenge in genomics, hampered by a high rate of false positive predictions^82^. Sequence-to-function models employing deep neural networks have emerged as promising methods to overcome this limitation^5,8–10^. The usage of chromatin accessibility data to train these models has been shown to be especially powerful to model enhancers, as differential accessibility has shown to be a key, and to our knowledge the most reliable, indicator of cell type-specific enhancer activity^22–24,29,31^. To make the training and downstream analysis of such scATAC-seq based models more robust and user-friendly, we developed CREsted. CREsted is a software package that enables training multi-class enhancer models across entire biological systems, obtaining enhancer codes, and designing enhancers with strong cell type-specificity.

By applying CREsted to a range of biological systems, we showed that CREsted can identify TFBS of key regulators to capture cell type-specific enhancer codes. We validated that these predicted TFBS overlap with functionally validated binding sites in human PBMC by comparing them against ChIP-seq peaks for a set of TFs active in various cell types. Furthermore, using single-cell multiomics data, we linked TFBS to potential TFs in the mouse motor cortex, whose expression correlates with the number of identified sites in cell type-specific regions. Most cell type-specific enhancers consist of clusters of TFBS. Whether these clusters form a rigid (i.e., “enhanceosome” model) or soft (i.e., “billboard” model) syntax is still an open question^83,84^. In this context, we studied an archetypal example of the enhanceosome model, the IFN-β enhancer^52,53^. CREsted was able to identify most of the validated TFBS in this enhancer region. Thus, while we do not yet answer this open question, we demonstrate that CREsted models can be used to investigate both the soft and the rigid syntax propositions. Further scrutiny of CREsted logic is therefore required. Besides exploring enhancer syntax, we highlight that in the context of evolutionary studies, CREsted can be useful to identify orthologous cell types and enhancers^29^. We showed that our mouse cortex model, DeepBICCN2, could be used to accurately predict accessibility in gene loci of orthologous chicken neuron subtypes even though they were entirely unseen by the model during training. Lastly, we explored a strategy, analogous to our previous work on comparing cross-species enhancer logic^29^, to compare cell states between tumor biopsies and cancer cell lines and identified a similar enhancer logic operating in the mesenchymal-like state in melanoma and glioblastoma cell lines that was partially recapitulated in the patient biopsies.

Next to modeling genomic enhancers, CREsted can be used to design synthetic enhancers. For this purpose, CREsted is used as a guiding oracle to optimize DNA sequences towards cell type-specificity for a desired class or set of classes. To design enhancers that are specific for selected cell types in the context of a whole organism we trained DeepZebrafish, a whole-organism CRested model trained on a zebrafish developmental scATAC-seq atlas^80^. To our knowledge, this is the first model capable of predicting cell type-specific accessibility across an entire organism, with the additional capability of maintaining this predictive power throughout developmental progression. Moreover, DeepZebrafish illustrates that CREsted scales to large datasets, here with 639 classes. We have previously shown this capacity with a full mouse brain model, DeepMouseBrain3, containing 221 classes^29^. By making use of the L2 distance we were able to design enhancers for somatic muscle, cardiac muscle, and endothelial cells with predicted accessibility within the range of genomic enhancers. These single cell type synthetic enhancers had a high success rate with all somatic and cardiac muscle enhancers, and one out of three endothelial enhancers being specifically active in the target cell type. This is especially remarkable given the high similarity of the enhancer codes of cardiac and somatic muscle cells (Fig. 6o). Previous work by Gosai *et al*.^32^ also validated synthetically designed sequences in zebrafish, targeting neurons and the liver. DeepZebrafish adds scale and resolution to these experiments, as we present the possibility of designing enhancers at cell type-specific resolution for the entire organism during development.

As a follow-up of our earlier work in the Drosophila brain^31^, we designed dual cell type enhancers (somatic and cardiac muscle), with overall more mosaic reporter expression. While designing these dual cell type enhancers, we designed multiple different sets that were suboptimal for one of the two cell types with the intent of modulating the level of GFP expression. We did observe an effect of the predicted level of expression, in the sense that enhancers that were predicted to have a higher level in somatic muscle cells compared to cardiac muscle cells, indeed labeled more somatic muscle cells compared to enhancers that were predicted to have a higher level of expression in cardiac muscle compared to somatic muscle cells. However, enhancers that were predicted to have equal levels of expression in both cell types labeled fewer somatic muscle cells in fewer embryos, compared to the enhancers that were designed to be stronger in somatic muscle cells (vis-à-vis cardiac muscle cells) even though the predicted level of somatic muscle cell chromatin accessibility of both enhancer sets is equal. This was an unexpected finding that warrants further investigation, as precise control over cargo expression levels in enhancer reporter assays holds significant relevance for disease-related gene therapy applications^85^. Of note, full quantification of expression levels may require the use of stable transgenics. In contrast, in our experiments, the observed reporter expression results from both the transient plasmid and the integrated enhancer.

CREsted is positioned within the rapidly expanding domain of enhancer modeling tools. One aspect of this domain aims at a complete understanding of gene regulation at a large scale, by predicting bulk or single-cell RNA-seq data and other genomic assays for large input sequences (hundreds of kilobases long). Models like Basenji^5^ and Enformer^10^ have pioneered this field, followed by successors like Borzoi^15^, Scooby^25^ and Decima^26^. While these models have shown to be powerful and can provide a global model of gene regulation across a wide variety of tissues, they may be overly complex for the task of decoding local enhancer regions. Furthermore, these large models are not optimized for enhancer design. Therefore, another direction entails local enhancer modeling, optimized for scrutinization of enhancer logic^9,13,14,21,28,30^. Complete software tools for modeling, analyzing and designing enhancers, however, are not as readily available. gReLU^35^ is to our knowledge the only comprehensive package providing all these options. Compared to gReLU, CREsted provides similar features, with additional peak normalization options and extensive enhancer code analyses. gReLU has a larger emphasis on the incorporation and utilization of the previously described large context models. We showed that multi-output task-specific models from CREsted outperform those trained in gReLU for a mouse motor cortex dataset. Other comprehensive packages, like tangermeme^46^, EUGENe^34^ and Selene^36^, either do not provide training or design options. Unlike ChromBPNet^8^ and scPrinter^27^, we decided not to pursue footprinting analysis, and to predict accessibility aggregated at a region level instead of base-pair level. This alleviated the need for Tn5 bias correction, while maintaining on-par enhancer code learning^8^. For example, in previous work, we have seen that results from both approaches result in very similar nucleotide contribution scores^29,31^. Moreover, we have extensively shown that CREsted infers TFBS with strong precision and recall through their contribution scores, both here and in previous studies^31^. We also validate CREsted models on large and complex scATAC-seq datasets, both here and in previous work^29^, an aspect that is often lacking in the aforementioned enhancer modeling packages.

CREsted models are multi-output, while footprinting models are trained on single scATAC tracks. The choice for a multi-output model has the benefit that it easily scales to large scATAC datasets comprising many cell types, as shown for DeepZebrafish, while training and evaluating many single-output models in parallel for such a large atlas is impractical. For our human cancer DeepCCL, we also observed that it outperformed a combined ChromBPNet model trained on separate cell lines. This could indicate that using the contrast of other cell types improves model performance, as has been observed in a previous study^86^. However, we have seen here and in past studies that multi-output models tend to misuse positive features, or motifs, of another class to negatively influence predictions for a given class. Such motifs are then represented with a negative contribution, even though they represent activator TFs expressed in other cell types and not repressor TFs as would be suggested by the attribution score. For example, in Fig. 2G, the pattern analysis suggested that DeepBICCN2 uses the CAGCTC E-box to not only correctly positively identify interneurons, but also to negatively impact excitatory neuron predictions. We did not find potential TF candidates that follow this pattern, suggesting that this may be a shortcut the model uses to differentiate global neuron types. Such issues should theoretically not be present in single-output models. Regardless, multi-output models can still identify repressive chromatin factors^31^, and using the global enhancer code analyses that CREsted provides can help filter out these potential false negatively contributing factors to the accessibility of a cell type.

Finally, we investigated the option of transfer learning our enhancer models from Borzoi. We showed that this technique results in near-identical results as training from scratch and outperforms predictions from the base Borzoi model. Since the transfer learning was done with the train, validation, and test splits as the DeepBICCN2 model, it was not aligned with the original data split used to train Borzoi, which could result in artificial inflation of model performance metrics through data leakage. However, transfer learning on the mouse motor cortex regions using the original Borzoi data split resulted in near-equivalent performance (Fig. S12b). Although transfer learning from a pre-trained model could have a larger benefit on datasets that are harder to model, e.g. due to a small amount of informative sequences to train on, this overall indicates that there is limited benefit in transfer learning from large pre-trained models for enhancer modeling and could potentially suggest that both approaches reach a performance plateau dependent on the data quality. For additional insights, extending these models to other genomic assays could be a relevant future research direction.

In summary, we present CREsted, a user-friendly software package integrated into *scverse*^37^ aimed at modeling, analyzing and designing enhancers from scATAC-seq data. We validated the package across multiple biological systems and species. The CREsted source code and all models we developed are freely available. The CREsted model repository also contains all the legacy models from our previous studies, including human, mouse and chicken brain^22,29^, mouse liver^30^, fly brain^21^ and human melanoma models^13,69^.

## Methods

### CREsted workflow

CREsted (RRID:SCR_026617) consists of four main modules: data preprocessing, model training, enhancer code analysis, and enhancer design. It is integrated into the *scverse*^37^ framework. Links to the internal tools used, the CREsted code and tutorials are found at https://crested.readthedocs.io and https://github.com/aertslab/CREsted.

#### Data preprocessing

CREsted preprocessing has two main modes. One is based on modeling peak heights from scATAC-seq tracks across cell types, the other uses topic modeling from pycisTopic^39,41^ (RRID: SCR_026618).

##### Peak height preprocessing

In the first mode, we use predetermined scATAC-seq tracks (BigWigs) per cell type. Here, we assume that tracks are already CPM-normalized. Then, using a predetermined consensus peak BED file, we obtain an aggregated peak height scalar per peak region using pybigtools^87^ (RRID:SCR_026627). The peak width is variable and selected by the user. We also ensure that peaks are not located outside their chromosomes after resizing. In the default setting, following an approach introduced by ChromBPNet^8^ (RRID:SCR_024806), we use peaks of width 2,114 bp, and calculate an aggregate peak height scalar for the center 1,000 bp. This can be done by either taking the max, mean, sum, or the logarithm of the sum of the cut sites or coverage inside the peak. Finally, we end up with a CxP matrix containing peak scalar values per cell type, with C the number of cell types and P number of peaks, and store that into an *anndata* object^37,88^ (RRID:SCR_018209).

##### Peak normalization

We rely on comparing peak heights across cell types, so we need to ensure those values are scaled appropriately. CPM-normalized tracks are only comparable if two tracks have a similar number of peaks, but when one of them has more peaks than the other, values will be lower overall because of the larger total amount of fragments used in to normalize the track (fig. S1A). We therefore apply a min-max normalization, by looking at the top peaks per cell type (configurable by user, default top 1%), filtering out peaks that are non-specific (Gini index less than the mean of Gini indices of all peaks minus one standard deviation) with the aim of only retaining strong, housekeeping constitutive peaks. We then calculate the average peak height of the strongest peaks per cell type, and obtain scalars per cell type which result in having the same average value per cell type. Finally, all peak values per cell type from the *anndata* object are updated by these cell type-specific peak scalars.

##### Topic modeling preprocessing

In the second mode, the outputs of pycisTopic^41^ are used to train a model that predicts topic probabilities for a given sequence. Specifically, binarized region-topic probabilities are used as the target class for each region in order to train a multi-label classification model. Compared to peak regression, topic classification requires less preprocessing since the data does not need to be normalized, the selected regions already include meaningful information captured by topic modeling, and consensus peaks are created on the fly from the provided topic regions. The width of the regions can be changed, but it is usually kept at the provided output width of pycisTopic which is 500 bp. Following a similar approach, there is also the ability to train regression models that use the extracted region-topic probabilities as target values.

##### Dataset splitting

In both modes we split the regions and their target values into a training, validation and test set either based on a manually defined chromosomal split or by randomly shuffling regions following a predefined percentage split.

#### Model training

For training CREsted models, there are also two main modes: regression and classification modeling. The regressions approach models peak scalars across cell types or topic probabilities across topics. The classification models determine the probability of a region belonging to a topic or a set of topics. We also provide the option of transfer learning from large pre-trained models, more specifically Enformer^10^ (RRID:SCR_024805) and Borzoi^15^ (RRID:SCR_026619). We use Keras 3.0 (RRID:SCR_026159) as a model training framework, which allows for both the TensorFlow (RRID:SCR_016345) and PyTorch (RRID:SCR_018536) backend.

##### Regression models

Regression models are multi-output and predict for a set of cell types or topic a scalar value. To train such models, we implemented as default loss function the sum of the cosine similarity between the predicted output vector and the ground truth peak height vector (both with length equal to the number of classes) and the MSE between the log-transformed predicted and ground truth scalars. To balance both parts of the loss function, we provide the option to dynamically scale the cosine similarity by the magnitude of the MSE. We provide other loss functions too, such as regular cosine similarity, Poisson loss and multinomial loss. We use the Adam optimizer with default learning rate set to 1e-3. We decay the learning rate by a factor of 4 after 5 (default) epochs without a decrease in validation loss and provide early stopping after 10 (default) epochs without a decrease in validation loss.

##### Fine-tuning to cell type-specific regions

We recommend the strategy of fine-tuning a peak regression model to a set of cell type-specific peaks. We define these peaks by calculating Gini indices of the target peak scalar vector of all peaks, and only retaining the regions with a Gini index above the mean of all Gini indices plus one standard deviation. Then, using the same regression modeling strategy, we train a regression model starting from the weights of a pre-trained model on all consensus peaks using a manually-defined smaller learning rate.

##### Classification models

Classification models are multi-label and predict probabilities for a set of topics^13,21,29,30^. We use the binary cross-entropy loss function to train these models. We use the Adam optimizer with default learning rate set to 1e-3. We decay the learning rate by a factor of 4 after 5 (default) epochs without a decrease in validation loss and provide early stopping after 10 (default) epochs without a decrease in validation loss.

##### Model architectures

For training enhancer models from scratch, there are multiple model architectures to choose from, both in the regression and classification approach. In regression models, the default model is a dilated CNN inspired by the ChromBPNet model architecture^8^, and there is also the option of using a Basenji-like architecture^5^ and a simple CNN. The topic models have as default the simple CNN model^29^, inspired by Basset^6^, and can also use a CNN + LSTM architecture^13,21^.

##### Transfer learning from pre-trained models

CREsted includes two large pre-trained models, Enformer^10^ and Borzoi^15^. These models both predict chromatin accessibility, DNA binding, and gene expression for many biological samples across a large genomic range, taking 192k bp or 512k bp input sequences to predict 114k bp and 192k bp of binned signal, respectively. To adjust these track coverage-predicting models for use with transfer learning on individual peak regions, we change them in two ways: shrinking the model’s input size to the size of a single region, and replacing the final layer to predict a vector of peak heights per region.

As both models bin to 128 bp internally, their new input size must be a multiple of 128 bp. We shrink these large models’ input size to 2,048 bp to approximate the *DilatedCNN* architecture input size. To change the model output from multiple-bin predictions to predicting a single vector of peak heights, we first disable the cropping layer and remove the final class-specific head layer. To replace the final layer, we instead flatten the model’s output embeddings and feed them to a single Dense layer with a Softplus activation, predicting the number of cell types desired. This results in a peak regression model benefitting from the weights of the original large pre-trained model.

We follow the default peak regression approach to train these models, except with lower batch size of 32 and a lower recommended learning rate of 5e-5 for first-round finetuning on the full peak set and 1e-5 for second-round finetuning on cell type-specific peaks respectively.

#### Enhancer code analysis

To obtain insights into underlying enhancer codes of model classes, we use nucleotide-level contribution scores.

##### Contribution score calculations

We calculate contribution scores for a sequence using gradient-based methods, which involve computing the gradient of the output with respect to the input. This highlights areas that the model focuses on. We implemented functions based on approaches used in the tfomics package^43^ to allow for a variety of options. To enhance the stability of these contribution scores, we provide integrated gradients and expected integrated gradients.

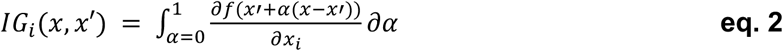

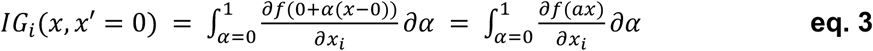

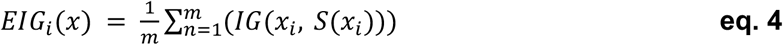

With:

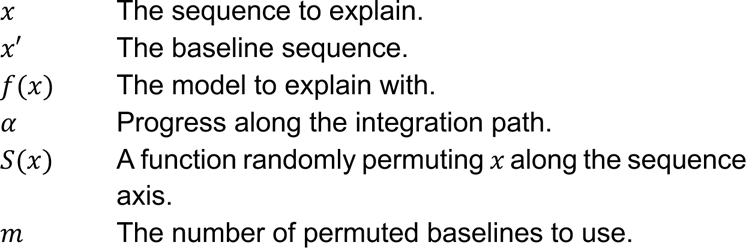

Integrated gradients^89^ (*IG*, eq. 2) calculate gradients for all sequences on the path from the baseline to the sequence to explain, and integrate over the resulting path of gradients. In our implementation, we use a zeroed-out sequence as a baseline (eq. 3), and approximate this integration by calculating the Riemann integral over gradients for 26 steps (*α*), including the baseline and final sequence. Expected integrated gradients^90^ (*EIG*, eq. 4), inspired by expected gradients^91^, replace the zeroed-out baseline by sampling the sequence space, randomly permuting the input sequence. This prevents the need for calculating gradients of fully zeroed-out sequences, which are normally out-of-distribution. CREsted permutes the input sequence 25 times (*m*) by default, averaging over the resulting integrated gradients calculated with respect to each baseline. By integrating information over several interpolations from a baseline, these techniques provide a more robust understanding of the model’s focus areas compared to simple gradient-based methods. This approach helps mitigate the noise and variability that can occur with single gradient calculations, offering a clearer and more reliable explanation of the model’s decisions.

We also provide the option to use in-silico mutagenesis (ISM) to further analyze the impact of mutations by generating mutated sequences and comparing them to the original, revealing how specific mutations affect the model’s output.

##### TF-MoDISco analysis

To obtain the most important patterns per cell type or topic, we identify frequently occurring motifs from the contribution scores of the most specific regions per cell type. To obtain those, we filter regions based on their peak specificity, their prediction specificity or the combination of both. We then take the top regions per cell type, and calculate contribution scores for the relevant class. These regions and contribution scores are used as input for tfmodisco-lite^44,45^ (RRID:SCR_024811), which we integrated into CREsted. We provide a motif database which is matched to the found motifs in a report.

##### Pattern clustering

To match the identified sequence patterns from multiple TF-MoDISco results, we implemented pattern clustering functionality. We read all patterns per cell type, and calculate their Tomtom similarity score with tangermeme^46^ (RRID:SCR_026620). We then group together patterns with a similarity score above a certain threshold, represented by the pattern with the highest information content. Single instance patterns with an information content below a defined threshold are discarded.

##### Pattern to TF matching

When the scATAC-seq data is part of a multiome experiment containing paired scRNA-seq data, we provide the option of automatically matching motifs to TF candidates. We do this by matching the importance vector of the motif over all cell types, to the average expression vector of a TF over all cell types, as was done in previous work^29^.

#### Enhancer design

##### Seed sequence generation

The enhancer design methods in CREsted are inspired by Taskiran *et al.* 2024^31^. Random DNA sequences of given length are generated that follow the GC content of genomic sequences. To this end, the fraction of each nucleotide of genomic sequences is calculated for each position and nucleotides are sampled according to this position-dependent distribution.

##### Optimization function

CREsted allows users to define their own optimization function. This function will be used during each iteration of the enhancer design process. CREsted ships with two types of optimization functions. The default function optimizes a weighted difference between a target class of interest and all other classes (eq. 5).

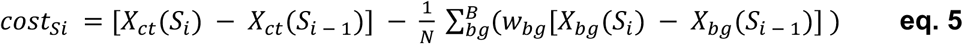

With:

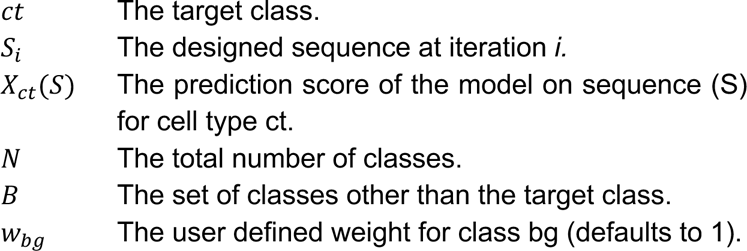

Another option is to use a function that optimizes the Euclidean distance between a predicted vector of chromatin accessibility to a user defined target one (see eq. 1 results section).

##### In silico evolution

A set of seed sequences is optimized by making all possible nucleotide substitutions for each sequence and selecting the most optimal substitution according to an optimization function. This process is repeated for a user-defined number of iterations^31^.

##### Motif embedding

A set of seed sequences is optimized by sequentially placing a set of user defined transcription factor binding sites in each sequence and selecting the most optimal location according to an optimization function, while ensuring that the newly placed binding site does not overlap previously placed binding sites^31^.

### Enhancer models

#### Model training

##### CREsted mouse cortex peak regression model, DeepBICCN2

We trained a multiclass regression model in CREsted on the mouse cortex dataset from Zemke *et al.*^47^ using the default peak regression configuration. Raw data from this dataset was obtained from GSE229169 and was processed as described in Johansen & Kempynck *et al.*^22^ to obtain cut-site BigWigs. The counts of the cut sites inside the center 1,000 bp of peaks were used as target values. Peaks were split into a training, validation and test set per chromosome (chr8 and chr10 in validation, chr9 and chr18 in test, remaining in train) such that they followed an approximate 80%-10%-10% distribution and they were resized to be of 2,114 bp length. We normalized peaks using the top 3% of peaks per cell type. Peaks were augmented by having stochastic 3 bp sequence shifts, and by reverse complementing all regions. We trained the model using the *DilatedCNN* model architecture, with as loss function the *CosineMSELogLoss* function (sum of the cosine similarity and the MSE on the log-transformed peak heights). The loss function was scaled dynamically. Adam optimizer was used with a learning rate of 1e-3. We pre-trained the model on 440,993 consensus peaks with a batch size of 256, and fine-tuned it to 73,326 cell type-specific regions obtained by filtering using the *crested.pp.filter_regions_on_specificity* function with a smaller batch size of 64 using a lower learning rate (1e-4). We had 440,993 peaks in our training set, 56,064 peaks in our validation set and 49,936 in our test set for the base model. For the fine-tuned model, we had 73,326 peaks in our training set, 9,951 in our validation set, and 8,198 in our test set. The whole training and evaluation procedures can be found in the *model_training_and_eval.ipynb* notebook.

##### gReLU mouse cortex peak regression model

We trained a multiclass regression model with the gReLU package^35^ (v. 1.0.3; RRID:SCR_026621), following the *3_train.ipynb* tutorial notebook. We provided the same set of cut-site BigWigs and consensus peaks from the Zemke *et al.* dataset^47^ used for training the CREsted peak regression model, and used the same train-validation-test chromosome split. Since we peak-scaled the peak heights across cell types in the CREsted model, we manually multiplied peak heights per cell type in gReLU with the same scalars. We also trained a larger gReLU regression model with 22M parameters instead of 6M, by doubling the number of convolutional filters (1024) from the original (512).

##### CREsted human PBMC peak regression model, DeepPBMC

We trained a multiclass regression model in CREsted on the human PBMC dataset from De Rop *et al.*^49^ using the default peak regression configuration. We downloaded coverage BigWigs from https://ucsctracks.aertslab.org/papers/scatac_benchmark/. The average coverage inside the center 1,000 bp of peaks were used as target values. Peaks were split into a training, validation and test set per chromosome (chr8 and chr10 in validation, chr9 and chr18 in test, remaining in train) and they were resized to be of 2,114 bp length. We normalized peaks using the top 3% of peaks per cell type. Peaks were augmented by having stochastic 3 bp sequence shifts, and by reverse complementing all regions. We trained the model using the *DilatedCNN* model architecture, with as loss function the *CosineMSELogLoss* function (sum of the cosine similarity and the MSE on the log-transformed peak heights). The loss function was scaled dynamically. Adam optimizer was used with a learning rate of 1e-3. We pre-trained the model on all 278,687 consensus peaks with a batch size of 128, and fine-tuned it to 51,644 cell type-specific regions obtained by filtering using the *filter_regions_on_specificity* function with a smaller batch size of 64 using a lower learning rate (1e-6). We had 278,687 peaks in our training set, 29,385 peaks in our validation set and 19,892 in our test set for the base model. For the fine-tuned model, we had 51,644 peaks in our training set, 6,070 in our validation set, and 4,303 in our test set.

##### CREsted cancer cell line peak regression model, DeepCCL

We trained a multiclass regression model on publicly available and newly generated data of eight cancer cell lines, including two DPCLs from ENCODE (HepG2, GM12878), three melanoma (MM029, MM099, MM001) and three GBM cell lines (A172, M059J, LN229). A detailed overview of the data generation and processing is included in the section Human cancer cell lines. Briefly, coverage BigWigs were generated for each cell line and peaks were called using MACS3. BigWig files of replicates were averaged using WiggleTools and peaks were merged into a consensus set using the *get_consensus_peaks* of pycisTopic^41^. These were then used to train a multiclass regression model on the mean coverage inside the center 1,000 bp of peaks. Peaks were split into a training, validation and test set per chromosome (chr7, chr8, chr9 and chr10 in validation, chr5 and chr6 in test, remaining in train) to follow a 70/20/10 distribution and were resized to a length of 2,114 bp. We normalized peaks using the top 3% of peaks per cell line. Peaks were augmented by having stochastic 5 bp sequence shifts, and by reverse complementing all regions. The *DilatedCNN* model architecture was used with Swish as an activation function and the filter size and dropout of the first convolutional layer was set to 11 and 0.2, respectively. The model was trained using the *PoissonLoss* over log-transformed values and the Lion optimizer with a learning rate of 5e-5 and a weight decay of 1e-2. We pre-trained the model on all 285,790 consensus peaks with a batch size of 512, and fine-tuned it to 140,761 cell-type specific regions with a batch size of 64 and a lower learning rate of 5e-6 and weight decay of 1e-1. Cell-type specific regions were obtained by calculating the coefficient of variation (CV) for the coverage across all regions and selecting the ones that have a CV above the median CV (equal to 0.9).

##### ChromBPNet cancer cell line model

We trained eight separate ChromBPNet^8^ models (i.e., one for each cell line) using the instructions provided at the corresponding GitHub page. We provided the same set of called peaks for each cell line and used the same train-validation-test chromosome split. The BAM files of replicates were merged as advised in the ChromBPNet tutorial. We evaluated the trained models using the obtained reports to confirm the effectiveness of the chosen hyperparameters. After training, we ported the models in CREsted by keeping only the count output head and integrating all eight models into one ensemble. Finally, to enhance the cell-type specific performance of the model and make the comparison fair, we fine-tuned the ChromBPNet ensemble to cell-type specific regions in a similar way to our DeepCCL training regime. We compared the performance of the ChromBPNet ensemble to the DeepCCL model by evaluating the Pearson correlation between log-transformed ground truth peak heights and model predictions (48,268 and 27,020 test set samples for the base and cell type-specific comparison, respectively) for each class (n = 8 classes).

##### CREsted glioma biopsy topic model, DeepGlioma

BAM files were downloaded from the European Genome-phenome Archive repository, under EGAS00001003845 and EGAD00001005314, and were processed into fragment files to be used as an input for pycisTopic^41^. First, pseudobulk ATAC-seq profiles were generated based on sample IDs and MACS2 (with a q-value equal to 0.001) was used to call peaks per pseudobulk. The pseudobulk peaks were then merged into a set of consensus regions with a size of 500 bp and were used for quality control and downstream analysis. Barcodes with less than 1,000 unique fragments, a TSS-enrichment below 5, and that were recognized as doublets were removed. This resulted in a cisTopic object of 20,316 cells and 429,016 regions. 30 topics were selected for topic modeling based on topic metrics. Sample batch effects were corrected with Harmony^92^. Visualization of the harmony-integrated tSNE after Leiden clustering (with a resolution of 0.4) demonstrated patient-specific clusters and smaller shared ones. After imputing region accessibility and inferring gene activity, we manually annotated the 14 identified clusters. We then reran the same pycisTopic workflow after excluding all the clearly identified healthy cells. This lead to a new cisTopic object of 14,275 cells and 311,118 regions. Similarly, we selected 25 topics based on topic metrics and corrected batch effects using Harmony^92^. Topic regions with the highest probability per topic were extracted after performing Otsu thresholding. One topic (Topic 3) included only 21 regions and was excluded from the subsequent analysis. The resulting binarized topic regions of 24 topics were used to train a CREsted multilabel classification model on a total of 192,256 regions. Regions were split into a training, validation and test set per chromosome (chr8, chr9 and chr10 in validation, chr5 and chr6 in test, remaining in train) to follow a 75/15/10 distribution and were kept at the length of 500 bp. Training regions were augmented by having stochastic 50 bp sequence shifts, and by reverse complementing all regions. The *DeepTopicCNN* model architecture was used with default parameters and trained using the *BinaryCrossentropy* loss and a batch size of 128. The Adam optimizer was user during training with a learning rate of 1e-3. The model with the lowest validation loss, obtained at epoch 61, was selected and evaluated using the area under the receiver operating characteristic curve and the area under the PR-curve for each topic.

##### CREsted zebrafish peak regression model, DeepZebrafish

The scATAC-seq fragment file and cell-level metadata file (containing cell type and development timepoint annotations) was downloaded from the gene expression omnibus (GEO) using identifier: GSE243256. The data was pre-processed using SnapATAC2 v2.6.4^40^. Briefly, consensus peaks were called by first calling peaks per cell type-timepoint combination using the function *snap.tl.macs3* followed by snap.tl.merge_peaks using default parameters. Next, a cut site consensus peak count matrix was generated using the function *snap.pp.make_peak_matrix* setting the counting_strategy option to “insertion”, finally a normalized matrix containing counts per cell type-timepoint combination was made using the function aggregate_X using the “RPM” normalization method. The resulting matrix was used to train the CREsted peak regression model. Consensus peaks were split in a validation (chromosome 8 and 10; n=70,993), test (chromosome 9 and 18; n=77,948) and training set (all chromosomes other than 8, 9, 10 and 18; n=793,273) and resized to 2,114 bp lengths. Peaks were augmented by having stochastic 3 bp sequence shifts, and by reverse complementing all regions. The *DilatedCNN* model architecture was used, using 1,024 filters in the first convolutional layer and 1,024 filters in the dilated convolutional layers. And the model was trained using the *CosineMSELogLoss*, which was scaled dynamically, and a batch size of 128. The Adam optimizer was used during training with a learning rate of 1e-3. The model that resulted after 53 epochs was finetuned for 3 more epochs on specific peaks (n=89,637 train, n=7,521 test and n=7,518 validate) that were determined using the *crested.pp.filter_regions_on_specificity* function using the same parameters as before but a learning rate of 2e-6.

##### CREsted Borzoi transfer learning

We trained models based on the Borzoi architecture and weights^15^ by initializing the CREsted-ported Borzoi model (replicate 0) as a base, with its input size reduced to 2,048 bp, cropping layer disabled, and final head layers (Conv1D + Softplus) disregarded, which results in a (64, 1536) output embedding. For the frozen Borzoi models, this base model’s weights were then frozen. The Borzoi CNN models followed the same architecture, except that the trunk was cut off after the last pooling layer of the convolution tower of the Borzoi model, resulting in a (16, 1536) output embedding. The trunk model was then extended with a Flatten and Dense layer, followed by a softplus activation function, to predict 19 pseudobulk classes.

These models were trained on the Zemke *et al.* dataset^47^, following the same data and training protocol as DeepBICCN2, except for the 2,048 bp input size, a batch size of 32, and a lower learning rate of 5e-5 for full-model first-round finetuning and 1e-5 for second-round finetuning and frozen model finetuning. The ‘fine-tuned’ Borzoi model was fine-tuned first on the full consensus peak set (matching the ‘base’ DeepBICCN2 model in fig. 2d), then further fine-tuned into ‘double fine-tuned’ on the cell type-specific peak set (matching the ‘fine-tuned’ DeepBICCN2 model). The ‘all-peak’ and ‘specific-peak’ frozen Borzoi models were fine-tuned for one round only on either the full consensus peak set or the cell type-specific peak set, as a single Dense layer does not benefit from multiple rounds of fine-tuning.

As the training-validation-test split used for DeepBICCN2 was not the same as the original Borzoi split, we also trained and evaluated models following the Borzoi data split. The consensus regions were assigned to a Borzoi genome fold by starting with Borzoi’s ‘sequences_mouse.bed’, merged into fold contigs with a gap tolerance of 10 bp, then intersected with the consensus regions, keeping regions that had any overlap with the folds. Regions overlapping multiple folds were assigned to the first fold when sorted by genomic position. Any consensus region not assigned a fold (3118 regions, 0.5%) was disregarded. The models were otherwise trained identically to the previously trained models, assigning fold 3 as test set and fold 4 as validation set. This data split resulted in 412,252 training, 72,747 validation and 58,876 test regions for the full dataset, and 67,153 training, 12,974 validation and 10,886 test regions for the cell type-specific dataset. Evaluating these models’ performance on the test set showed similar performance to the original data split-trained models (fig. S12b).

For all models, the lowest validation loss epoch was kept as the final model. For the non-frozen models, this happened within 5 epochs. The frozen models trained slower, and were left to train for 30 epochs, with automatic learning rate lowering applied with a patience of 5, as is the CREsted default.

#### Model evaluation

##### CREsted and gReLU performance comparison

We compared performance metrics across models by evaluating the mean Pearson correlation between log-transformed ground truth peak heights and model predictions (49,936 and 8,198 test set samples for base and specific comparisons respectively) for each class (n=19 classes). We used independent two-sided Welch’s t-tests for all pairwise model comparisons, which account for potential unequal variances between groups. For each comparison, we tested whether the distributions of per-class correlation values differed significantly between models. Each correlation value corresponds to a distinct class and is computed using non-overlapping data points per model (i.e., predictions from different models on the same evaluation data). As the unit of analysis is the class-level correlation, and classes are evaluated only once per model, measurements were not repeated across models. We performed all possible pairwise comparisons between the selected models. For each, we report the t-statistic, degrees of freedom, and two-sided p-value. To correct for multiple hypothesis testing, we applied the Benjamini–Hochberg procedure for false discovery rate control (using method="fdr_bh" from statsmodels.stats.multitest.multipletests). All statistical tests were conducted in Python using scipy.stats.ttest_ind and statsmodels for multiple testing correction.

##### Gene locus predictions

We scored gene loci by means of a sliding window of 2,114 bp predictions, shifted 100 bp every step. We defined the gene locus as the genomic region from 50 kb upstream of the TSS to 25 kb downstream the 3’ UTR. For a given genomic position, the final predicted peak height is the average of all the overlapping predictions that position is part of. These predictions were made with *crested.tl.score_gene_locus*. Visualizations were made with *crested.pl.hist.locus_scoring*.

##### CREsted predictions on validated mouse cortex enhancers

A set of 171 functionally validated, On-Target, mouse and human genomic enhancer regions in the mouse motor cortex were taken from a dataset provided by Ben-Simon *et al*.^23^. Models were assessed by evaluating their classification performance on this set. This was done by taking the target cell type and optional second target cell type, and setting those as ground truth labels per enhancer. Then, we scored all regions, and took the highest predicted class as a predicted label, both for predictions from CREsted models and for scATAC peak heights. Since the enhancers were all shorter than 2,114 bp and had variable lengths, we zero-padded the flanks to complete the sequence to 2,114 bp to use as input for CREsted models. For scATAC peaks, we took the exact values for the length of the enhancer, and zero-padded values to 1,000 bp if the enhancer was shorter as the CREsted models also predict a value over a 1,000 bp window.

We also checked specificity of the predictions by using the *crested.pp._utils._calc_proportion* function. By using the ground truth labels obtained in the classification step, we calculated precision and recall per class using different specificity thresholds, as was done in previous work^22^. For scoring Base Borzoi, we centered the 524,288 bp sequences on the enhancer regions, and used their genomic flankings. We then scored all regions for the relevant mouse cortex classes, and took as predicted values the center 32 bins, corresponding to 1024 bp. We then calculated specificity by only comparing the 19 relevant classes.

##### EBF1 degradation analysis in B-cells

To simulate the EBF1 degradation experiment^63^ and its effect on accessibility in B-cells around the *TCF3* gene locus, we identified all exact hits of CCCCTGGG, CCCTAGGG and TCCCTGGG to find potential EBF1 TFBS and replaced those by N’s, resulting in a zero-input. We rescored the gene locus with this modified genome with *crested.tl.score_gene_locus*, going 20 kb upstream and 5 kb downstream of the gene position.

##### CREsted predictions against topics derived from the cisTopic analysis of human Gliomas

We generated pseudobulk topic bigwig files by binarizing cell-topic distributions and assigning cells to specific topics. Pseudobulk profiles were then created using the *export_pseudobulk* function in pycisTopic^41^ for each topic. Additionally, we identified differentially accessible regions for each annotated cluster and selected the top 10,000 for downstream comparison. Using these regions, we compared DeepCCL predictions (as well as cell line accessibility) with topic accessibility. Specifically, we averaged the predictions and accessibility of GBM and melanoma MES-like cell lines, then applied Spearman correlation to identify potential biopsy topics that share a similar MES-like regulatory landscape. Similarly, we used the predictions/accessibility of LN229 to find candidate OPC/NPC-like topics.

##### CREsted predictions on validated zebrafish enhancers

A set of 100 tested enhancers (supplementary table 6 in Sun *et al.*^80^) were rescaled to 2,114 bp centered on the middle of each enhancer, 54 of these enhancers were active in at least one cell type and were scored using the DeepZebrafish model.

##### Comparisons of CREsted-trained models with base Borzoi

The CREsted Borzoi transfer learning models were compared with pre-computed BigWig pseudobulk signal tracks from Zu *et al*.^16^ (http://catlas.org/wholemousebrain/), with selected cell types corresponding to the Zemke *et al.*^47^ cell types (fig. S12b). Values from these BigWig tracks were read into CREsted using the cell type-specific region set from the DeepBICCN2 analysis (fig. 5a), taking the mean of the signal inside the 1000bp at the center of each region.

The base Borzoi predictions were calculated with the CREsted-included base Borzoi model, using the mouse head of replicate 0. To calculate the scores, a 524288 bp region from the mm10 genome was extracted surrounding each peak, padding with zeros where it fell outside chromosome boundaries. The core 32 bins (corresponding to 1024bp) of the prediction were then summed and saved for each of the corresponding classes in fig. 5e. Notably, reversing the transformations (taking mean of 32bp bins, soft-clipping, and scaling) applied to Borzoi’s training data to approximate the original fold-change scores worsened the correlations with the Zu *et al.* data.

The contribution scores were predicted using *crested.tl.contribution_scores()*, which uses expected integrated gradients as its default. For the base Borzoi model, the attributions were calculated for the annotated track at bin 3071, for a full 524288 bp sequence centered on the region to be evaluated.

#### Motif analysis

##### Mouse cortex motif analysis

To obtain top patterns per mouse cortex cell type, we used the top 2,000 most specific regions per cell type. We obtained these by averaging both the peak and prediction specificity. We then ran the standard CREsted enhancer code analysis pipeline, by calculating contribution scores per cell type through *crested.tl.contribution_scores_specific,* running *tfmodisco-lite* with *crested.tl.modisco.tfmodisco(window=1000, max_seqlets=20000)* and clustering patterns with *crested.tl.modisco.process_patterns*(*sim_threshold = 4, trim_ic_threshold = 0.025, discard_ic_threshold = 0.2)* and *crested.tl.modisco.create_pattern_matrix(normalize=False, pattern_parameter="seqlet_count_log")*. We then visualized the clustering with *crested.pl.patterns.clustermap_with_pwm_logos(importance_threshold=5,logo_height_fraction= 0.35,logo_y_padding=0.25)*.

##### Human PBMC motif analysis

To obtain top patterns per human PBMC cell type, we used the top 1,000 most specific regions per cell type. We obtained these by averaging both the peak and prediction specificity. We then ran the standard CREsted enhancer code analysis pipeline, by calculating contribution scores per cell type through *crested.tl.contribution_scores_specific,*running *tfmodisco-lite* with *crested.tl.modisco.tfmodisco(window=1000, max_seqlets=20000)* and clustering patterns with *crested.tl.modisco.process_patterns*(*sim_threshold = 4.25, trim_ic_threshold = 0.05, discard_ic_threshold = 0.2)* and *crested.tl.modisco.create_pattern_matrix(normalize=False, pattern_parameter="seqlet_count_log")*. We then visualized the clustering with *crested.pl.patterns.clustermap_with_pwm_logos(importance_threshold=4.5,logo_height_fractio n=0.4,logo_y_padding=0.25)*.

##### Human PBMC ChIP-seq comparison

We evaluated the identification of seqlets of PAX5, EBF1 and POU2F2 in B cells, GATA3, RUNX1 and ETS1 in CD4+ T cells and CEBPA and SPI1 in CD14+ monocytes by comparing them with ChIP-seq peaks. We downloaded ChIP-seq data, including peak files and source bigwigs, for PAX5 from ENCODE ENCFF827VVQ and ENCFF914QGY, for EBF1 from ENCODE ENCFF895MHN and ENCFF810XRY, for POU2F2 from ENCODE ENCFF803HIP and ENCFF934JFA. Additionally, we retrieved ChIP-seq data from ChIP-Atlas (https://chip-atlas.org/) for GATA3 (SRX4705120), RUNX1 (SRX1492212), ETS1 (SRX015825), CEBPA (SRX097095), and SPI1 (SRX4001818).

Then, we took the top 1,000 most specific regions per cell type from the motif analysis, and assigned motifs to TFs based on tomtom matches with our motif database^39^. We compared the individual instances, or seqlets, of these CREsted-identified TFBS to the called ChIP-seq peaks. We calculated precision and recall, where true positives (TP) indicate seqlets inside a ChIP peak, false positives (FP) seqlets that are not and false negatives (FN) ChIP peaks without a seqlet and thresholded the selection of seqlets based on their average contribution score to make a precision-recall curve.

##### Human PBMC UniBind comparison

We downloaded UniBind hits to obtain potential precise TFBS of the TFs we obtained ChIP-seq data from. For B cells, we downloaded hits for PAX5 and EBF1 from https://unibind.uio.no/factor/ENCSR000BHD.GM12878_female_B-cells_lymphoblastoid_cell_line.PAX5/ and https://unibind.uio.no/factor/ENCSR000DZQ.GM12878_female_B-cells_lymphoblastoid_cell_line.EBF1/ For CD4+ T cells, we downloaded hits for GATA3, RUNX1 and ETS1 from https://unibind.uio.no/factor/GSE76181.Jurkat_T-cells.GATA3/, https://unibind.uio.no/factor/GSE76181.Jurkat_T-cells.RUNX1/ and https://unibind.uio.no/factor/EXP000299.Jurkat_E6_1_T-cells.ETS1/. For CD14+ monocytes, we downloaded hits for CEBPA and SPI1 from https://unibind.uio.no/factor/EXP000946.U937_adult_acute_monocytic_leukemia.CEBPA/ and https://unibind.uio.no/factor/EXP047756.MDMmonocyte_derived_macrophages.SPI1/.

We checked if at those genomic positions DeepPBMC identified a seqlet. We obtained seqlets from sequences through the *recursive_seqlets* function with a P-value parameter of 0.05 in *tangermeme*^46^. We then calculated overlap between identified seqlets in a region and UniBind hits, and called a match when the overlap was more than 50%. We did this both for UniBind hits inside the top 1,000 regions per cell type, and for hits inside all consensus peaks.

##### Glioblastoma motif analysis

For the DeepCCL model, we used the top 2,000 most specific regions per cell line to obtain their top patterns. We obtained these by averaging both the peak and the prediction specificity. We then ran the standard CREsted enhancer code analysis pipeline, by calculating contribution scores per cell line through *tfmodisco_calculate_and_save_contribution_scores* and running tfmodisco-lite with *crested.tl.modisco.tfmodisco(window=1000, max_seqlets=20000)*. The same process was performed for the base and final DeepCCL model and their identified patterns were clustered with *crested.tl.modisco.process_patterns(sim_threshold=3.5, trim_ic_threshold = 0.1, discard_ic_threshold = 0.2)* and *crested.tl.modisco.create_pattern_matrix(normalize=False, pattern_parameter="seqlet_count")*. In the resulting pattern matrix, the seqlet counts of the A172/M059J and the MM029/MM099 cell lines were averaged and log-transformed, while keeping the original sign of the contribution scores. Not meaningful seqlets and seqlets with zero count in both GBM and melanoma cell lines were removed, and the remaining clusters were manually annotated for plotting.

For the DeepGlioma model, we selected 7 relevant topics (Topics 8, 21, 25, 20, 14, 18, 19) to focus our enhancer code analysis. We selected the top 2,000 regions per topic and ran the standard CREsted enhancer code analysis pipeline, by calculating contribution scores per topic through tfmodisco_calculate_and_save_contribution_scores and running tfmodisco-lite with *crested.tl.modisco.tfmodisco(window=500, max_seqlets=20000)*.

For the combined analysis, we selected the top 1,000 most specific regions per cell line by averaging the peak and the prediction specificity, and calculated the contribution scores of each DeepCCL output for each cell line through *tfmodisco_calculate_and_save_contribution_scores.* Similarly, we selected the top 1,000 regions per topic and obtained the contribution scores for the corresponding topic outputs of DeepGlioma. In addition, we calculated the contribution scores of all DeepCCL output cell lines for each selected topic. Then we calculated the Spearman correlation of the obtained contribution scores for each region, by taking the middle 500 bp where necessary (i.e., when comparing the DeepCCL and DeepGlioma contributions). The correlations of the top 1,000 regions were aggregated by calculating the median to obtain a similarity score between all cell lines and biopsy topics on a nucleotide contribution level.

Finally, the previously identified patterns of DeepCCL and DeepGlioma were clustered together using *crested.tl.modisco.process_patterns(sim_threshold = 3.5, trim_ic_threshold = 0.1, discard_ic_threshold = 0.2)* and *crested.tl.modisco.create_pattern_matrix(normalize = False, pattern_parameter = "seqlet_count_log").* The resulting clustering was visualized with *crested.pl.patterns.clustermap_with_pwm_logos(importance_threshold = 5.0)* and subsetting to the classes that we wanted to compare (e.g., MES-like cell lines and topics, LN229 and OPC/NPC-like topics).

##### Zebrafish motif analysis

To obtain patterns for the enhancers designed for either cardiac muscle or somatic muscle or both contribution scores were calculated for all designed enhancers for the classes: “Slow muscle cells”, “Fast muscle cells” and “Cardiac muscle” of development stage 72 hpf using the function *tfmodisco_calculate_and_save_contribution_scores_sequences* and method *integrated_grad*. Next, *tfmodisco-lite* was run using the *crested.tl.tfmodisco* using default parameters. Patterns were clustered using the function *crested.tl.modisco.process_patterns* and parameters sim_threshold=4.25, trim_ic_threshold=0.05, and discard_ic_threshold=0.15 and visualized in a heatmap using the function *clustermap_with_pwm_logos* using parameter importance_threshold=4. To obtain patterns for the enhancers designed for endothelial cells, contribution scores were calculated for all designed enhancers for the class “Endothelial” of development stage 72 hpf using the function *tfmodisco_calculate_and_save_contribution_scores_sequences* and method *integrated_grad*. Next, *tfmodisco-lite* was run using the *crested.tl.tfmodisco* using default parameters.

#### Zebrafish enhancer design

##### Zebrafish cardiac and body muscle enhancer design

Cell type classes of any time point annotated as either: “Slow muscle cell”, “Slow muscle cells”, “Fast muscle cells”, “Heart”, “Heart field” or “Cardiac muscle” were considered and the average over all positive enhancers (see “CREsted predictions on validated zebrafish enhancers”) of the maximum prediction score over all classes was calculated (*avg_pos_prediction_score*). To design enhancers specific for cardiac muscle a cost function was defined that minimizes the euclidean distance between the models prediction score on the set of considered cell types and a target vector where classes corresponding to “Cardiac muscle” equals *avg_pos_prediction_score* and other cell types are zero. Similarly, for body muscle a target vector was used where classes corresponding to either “Slow muscle cell”, “Slow muscle cells” or “Fast muscle cells” equals *avg_pos_prediction_score* and the other cell types are zero. Three sets of enhancers were designed to be active in both cell types, with varying levels of prediction scores: cardiac_1_/body_1_ with equal prediction score in cardiac and body muscle cells, cardiac_1_/body_0.5_ where the prediction score in body muscle cells in half the prediction score in cardiac muscle cells and vice versa cardiac_0.5_/body_1_. For cardiac_1_/body_1_ a target vector was used where classes corresponding to “Cardiac muscle”, “Slow muscle cell”, “Slow muscle cells” or “Fast muscle cells” equals *avg_pos_prediction_score* and the other cell types are zero. For cardiac_1_/body_0.5_ a target vector was used where classes corresponding to “Cardiac muscle” equals *avg_pos_prediction_score* and classes corresponding to Slow muscle cell”, “Slow muscle cells” or “Fast muscle cells” equals *avg_pos_prediction_score* divided by 2. Vice versa for cardiac_0.5_/body_1_. Random DNA sequences were initialized with similar nucleotide content across the peak region as consensus peaks. For this the fraction of each nucleotide for each position in the peak was calculated on the test set.

Using this, for each set of designed enhancers 200 random DNA sequences were generated and were optimized using in silico evolution. See supplementary table S1.

##### Zebrafish endothelial enhancer design

The same approach as was used to design enhancers targeted for either cardiac or body muscle was used to design endothelial cells. For this class corresponding to “Notochord”, “Endothelial” or “Otic placode” were considered and a target vector where Endothelial was set to *avg_pos_prediction_score* and the others to zero was used. See table S1. Cancer cell line analysis

### Human cancer cell lines

#### Culture of human GBM cell lines

A172 cells (ATCC Cat# CRL-1620; RRID:CVCL_0131) were cultured in DMEM (ThermoFisher 11965-092) medium supplemented with 10% FBS (ThermoFisher 17479-633), and 1% penicillin/streptomycin (Life Technologies 15140122). MO59J cells (ATCC Cat# CRL-2366; RRID:CVCL_0400) were cultured in DMEM/F12 (ThermoFisher 11320-033) medium supplemented with 10% FBS, 1% penicillin/streptomycin and 0.05 mM non-essential amino acids (ThermoFisher 11140-068). LN229 cells (ATCC Cat# CRL-2611; RRID:CVCL_0393) were cultured in DMEM (ThermoFisher 11965-092) medium supplemented with 5% FBS and 1% penicillin/streptomycin. All cell lines were passaged twice per week with Trypsin-EDTA 0.05% (ThermoFisher 11580-626).

#### Bulk ATAC-seq

Cells were treated with Trypsin-EDTA 0.05% and washed with DPBS (ThermoFisher 14190-169). Cell viability and concentration were assessed by the LUNA-FL Dual Fluorescence Cell Counter. Per cell line, 80,000 cells were used to perform ATAC-seq. After centrifugation at 500 g for 5 min at 4°C, the supernatant was replaced with 50 µl ice cold ATAC-seq lysis buffer (10 mM Tris-HCl, 10 mM NaCl, 3 mM MgCl2, 0.1% IGEPAL CA-630, 0.1% Tween-20, 1% BSA and 0.01% Digitonin) and the cells were mixed by pipetting. After incubation for 5 min on ice, 1 ml of wash buffer (20 mM Tris-HCl, 20 mM NaCl, 6 mM MgCl2, 0.1% Tween-20, and 1% BSA) was added to the lysed cells and mixed gently by inverting 3 times. After centrifugation at 500 g for 5 min at 4°C, supernatant was removed and the nuclei were resuspended in 50 µl ATAC mix (10 mM Tris-HCl, 10% dimethylformamide, 5 mM MgCl2, 1x DPBS, 0.1% Tween-20, 0.01% Digitonin and 3.75 ng/µl Tn5 enzyme) and incubated for 1 hour at 37°C. Transposed DNA was purified using a QIAGEN MinElute purification column and eluted in 15 µl elution buffer (Qiagen). The ATAC-seq library was completed by amplifying the transposed DNA in a total volume of 50 µl PCR mix (15 µl purified DNA, 25 µl NEBNext High Fidelity PCR Master Mix (NEB), 2.5 µl FWD and 2.5 µl REV primer) with the following program: 72°C for 5 min, 98°C for 30 s, followed by 10 cycles of 98°C for 10 s, 63°C for 30 s, and 72°C for 1 min). The final libraries were subjected to 0.4x-1.2x doublesided Ampure purification and eluted in 20 µl elution buffer (Qiagen).

#### Sequencing

ATAC-seq libraries were sequenced on an Illumina NovaSeqX system using 51 cycles for read 1, (ATAC paired-end mate 1), 8 cycles for index 1 (sample index 1), 8 cycles for index 2 (sample index 2) and 51 cycles for read 2 (ATAC paired-end mate 2).

#### Data processing

FASTQ files for HepG2 (RRID:CVCL_0027) and GM12878 (RRID:CVCL_7526) were downloaded from ENCODE^59,70^ (https://www.encodeproject.org/, RRID:SCR_006793) under accession number ENCLB324GIU and ENCLB907YRF, respectively. The quality of these reads together with the newly generated ones for A172, M059, and LN229 was evaluated using FastQC (0.11.9, RRID:SCR_014583) and any adapters were trimmed using the fastq-mcf command from ea-utils (1.1.2.779, RRID:SCR_005553). Reads were mapped to the hg38 genome using Bowtie2^93^ (2.4.4, RRID:SCR_016368) and duplicates were removed using picard (2.27.1, RRID:SCR_006525). Peaks were called using MACS3^94^ (3.0.0b1, RRID:SCR_013291) and aggregate genome coverage was generated with the bamCoverage command of deepTools^95^ (3.5.0, RRID:SCR_016366). The coverage of replicates was averaged using WiggleTools (1.2.11, RRID:SCR_001170). For the melanoma cell lines, fragment files were downloaded from Gene Expression Omnibus (GEO) with accession number GSE210745. Pseudobulk ATAC-seq profiles were generated per cell line as shown in the SCENIC+ tutorial^39^ and peaks were called for each pseudobulk. From these, we selected MM001, MM029, and MM099 for our analysis. The resulting peaks from all eight cell lines (i.e., the melanoma, GBM, and ENCODE ones) were merged into a consensus set using the *get_consensus_peaks* function of pycisTopic^41^ and were used together with the eight coverage files to train the DeepCCL model. ChIP-seq peaks for JunB, cJun, and cFos in MM099 were downloaded from GEO using identifier GSE159965 and for ZEB1 through https://ucsctracks.aertslab.org/papers/enhancer_design/hg38/bb/tf_chip/. In addition, we retrieved ChIP-seq peaks for TEAD1 in MSTO cells and TEAD4 in SK-MEL-147 through ReMap^96^ with accession numbers GSE68170 and GSE94488, respectively.

### Enhancer reporter assays

#### *In vivo* validation of zebrafish enhancers

##### Enhancer-reporter plasmid

Synthetic enhancer DNA sequences were ordered from Twist Bioscience with constant flanking DNA sequences: 5’ end “ATATACCCTCTAGAGTCGAA” and 3’ end “GATTACCCTGTTATCCCTAA”. These flanking regions were used to PCR-amplify the sequences using primers containing overhangs that are homologous to the target plasmid.

Fwd: 5’ TTAGGGATAACAGGGTAATCGCGAATTGGGTACCGGGC 3’
Rev: 5’ CTTTCAACAAGCCCGAAAGATCTTCTGGAAGCCTCCAGTGAATT 3’

We used the KAPA HiFi HotStart ReadyMix (Roche) with primer and template concentrations of 300 µM and 0.1 ng/ul, respectively. PCR setup: 95C 3’, 20 cycles of 98C 20’’, 65C 15’’, 72C 15’’, and a final elongation step of 72C 2’. Next, the target plasmid, Tol2-ISceI- ZSP:EGFP;cryaa:mCherry-ISceI-Tol2 (addgene id #194518) was linearized by restriction digestion using BglII and XhoI (at 0.5 and 1 U/µl, respectively, in rCutSmart buffer, with 7 µg plasmid, 2 h incubation at 37°C), purified (NucleoSpin Gel & PCR cleanup kit, Macherey-Nagel) and combined with purified enhancer DNA fragments in a NEBuilder reaction (NEBuilder enzyme mix from New England Biolabs, 7 fmol linearized plasmid, 65 fmol DNA fragment, 45 min incubation at 50°C). Then, 2.5 µl of the reaction was transformed into 20 µl of Stellar chemically competent bacteria (thaw cells on ice for 15 min, add plasmid, keep 30 min on ice, 45 sec heat shock at 42°C, 5 min ice), which were incubated overnight at 37°C on carbenicillin plates. Single colonies were then grown in 10 ml LB medium, and plasmids were isolated using QIAprep Spin Miniprep Kit (Qiagen) and eluted in 2x25 µl. All plasmid preps were sequenced using Sanger sequencing to confirm correct insertion in the target plasmid without sequence alterations. Enhancers C1, B1, F1, G1 and G2 contain point mutations that should not affect enhancer activity according to the model prediction.

##### Egg microinjection

Plasmid DNA together with Tol2 messenger RNA were injected (at concentrations of 30 and 40 ng/µl, respectively) in one-cell stage zebrafish eggs (of the wild type AB strain) and they were grown at 28.5C until 48 hpf. Zebrafish with successful injection were selected based on red fluorescence in the eye on a Olympus SZX16 widefield fluorescence microscope. Fish were anesthetized with 0.02% tricaine (MS-222 Ethyl 3-aminobenzoate methanesulfonate, Sigma) and mounted in 1% low melting point agarose (invitrogen) on fluorodish (FD3510-100. world precision instruments). All zebrafish breeding was approved by the Ethical Committee for Animal Experimentation of the KU Leuven (ECD-000) and all experiments were performed on embryos younger than five days post fertilization.

##### Imaging

48 hpf zebrafish embryos were imaged for GFP fluorescence on a spinning disk confocal microscope consisting of a Nikon Ti2 body and Crest X-Light V2 spinning disk using a 10x Plan Apo Lambda lens with numerical aperture of 0.45 and 20x Plan Apo Lambda lens with numerical aperture of 0.8. The Lumencor Spectra III using the green channel was used to excite GFP fluorescence, using excitation filter Semrock FF01-378/474/554/635/735-25 and dichroic mirror FF409/493/573/652/759-Di01-25x36. Semrock FF01-515/30-25 was used as an emission filter. NIS element V6 software was used to control the microscope.

## Data and code availability

The CREsted package is available at https://github.com/aertslab/CREsted and https://crested.readthedocs.io and is stored at https://zenodo.org/records/15045960. All computational analyses for the main figures can be found in https://github.com/aertslab/CREsted-paper and are stored at https://zenodo.org/records/15111374. Analysis data required for the notebooks is available at https://resources.aertslab.org/CREsted/manuscript_data/. All CREsted models developed in this paper, and other legacy models, can be loaded through *crested.get_model*, or downloaded directly from https://resources.aertslab.org/CREsted/. All raw and processed sequencing data generated in this study have been deposited in NCBI’s Gene Expression Omnibus and are accessible through GEO Series accession number GSE292617. This includes the OmniATAC-seq data of three human glioblastoma cell lines, namely A172, M059J, and LN229. All the datasets, software, protocols, and lab materials used and/or generated in this study are listed in a Key Resource Table alongside their persistent identifiers in Supplementary Table S2.

## Authors’ contributions

Conceptualization: N.K., S.A.

Computational analysis: N.K., S.D.W, C.H.B., V.K., S.D.

Data collection, processing, and curation: N.K., S.D.W., V.K., S.D.

Experiments and sample preparation: S.D.W., V.B., K.S., V.C.

Resources: S.A., L.V.D.B.

Software implementation and testing: L.M., N.K., C.H.B., I.I.T., S.D.W., E.C.E., V.K., G.H.

Visualization: N.K., S.D.W., C.H.B., V.K.

Writing – original draft: N.K., S.D.W, C.H.B., V.K., S.A.

Writing – review & editing: N.K., S.D.W, C.H.B., V.K., S.A., E.C.E., L.M., S.D., V.B.

## Acknowledgments

We want to thank the members of the Lab of Computational Biology for discussions and feedback on the CREsted package and manuscript, in particular N. Hecker, H. Dickmänken, D. Abaffyová, K. Theunis, O. Sigalova, D. Daaboul, F. De Rop and C. Bravo González-Blas. We also thank early external users of the package for their feedback. We thank the staff of the VIB Data Core and the Flemish Supercomputing Center (Vlaams Supercomputer Centrum - VSC) for their support. An element of Fig. 4a was obtained from BioRender. Finally, we thank D. Koldere Vilain for the CREsted logo design.

## References

1. Davidson, E. H. *cis*-Regulatory Modules, and the Structure/Function Basis of Regulatory Logic. in The Regulatory Genome (ed. Davidson, E. H.) 31–86 (Academic Press, Burlington, 2006). doi:10.1016/B978-012088563-3.50020-1.

2. Zeitlinger, J. Seven myths of how transcription factors read the cis-regulatory code. Curr. Opin. Syst. Biol. 23, 22–31 (2020).

3. Zhou, J. & Troyanskaya, O. G. Predicting effects of noncoding variants with deep learning-based sequence model. Nat. Methods 12, 931–934 (2015).

4. Chen, K. M., Wong, A. K., Troyanskaya, O. G. & Zhou, J. A sequence-based global map of regulatory activity for deciphering human genetics. Nat. Genet. 54, 940–949 (2022).

5. Kelley, D. R. et al. Sequential regulatory activity prediction across chromosomes with convolutional neural networks. Genome Res. 28, 739–750 (2018).

6. Kelley, D. R., Snoek, J. & Rinn, J. L. Basset: learning the regulatory code of the accessible genome with deep convolutional neural networks. Genome Res. 26, 990–999 (2016).

7. Yuan, H. & Kelley, D. R. scBasset: sequence-based modeling of single-cell ATAC-seq using convolutional neural networks. Nat. Methods 19, 1088–1096 (2022).

8. Pampari, A. et al. ChromBPNet: bias factorized, base-resolution deep learning models of chromatin accessibility reveal cis-regulatory sequence syntax, transcription factor footprints and regulatory variants. 2024.12.25.630221 Preprint at 10.1101/2024.12.25.630221 (2024).

9. Avsec, Ž., et al. Base-resolution models of transcription-factor binding reveal soft motif syntax. Nat. Genet. 53, 354–366 (2021).

10. Avsec, Ž., et al. Effective gene expression prediction from sequence by integrating long-range interactions. Nat. Methods 18, 1196–1203 (2021).

11. Quang, D. & Xie, X. DanQ: a hybrid convolutional and recurrent deep neural network for quantifying the function of DNA sequences. Nucleic Acids Res. 44, e107 (2016).

12. De Winter, S., Konstantakos, V. & Aerts, S. Modelling and design of transcriptional enhancers. Nat. Rev. Bioeng. 1–16 (2025) doi:10.1038/s44222-025-00280-y.

13. Minnoye, L. et al. Cross-species analysis of enhancer logic using deep learning. Genome Res. gr.260844.120 (2020) doi:10.1101/gr.260844.120.

14. de Almeida, B. P., Reiter, F., Pagani, M. & Stark, A. DeepSTARR predicts enhancer activity from DNA sequence and enables the de novo design of synthetic enhancers. Nat. Genet. 54, 613–624 (2022).

15. Linder, J., Srivastava, D., Yuan, H., Agarwal, V. & Kelley, D. R. Predicting RNA-seq coverage from DNA sequence as a unifying model of gene regulation. Nat. Genet. 1–13 (2025) doi:10.1038/s41588-024-02053-6.

16. Zu, S. et al. Single-cell analysis of chromatin accessibility in the adult mouse brain. Nature 624, 378–389 (2023).

17. Li, Y. E. et al. A comparative atlas of single-cell chromatin accessibility in the human brain. Science 382, eadf7044 (2023).

18. Gao, Y. et al. Continuous cell type diversification throughout the embryonic and postnatal mouse visual cortex development. 2024.10.02.616246 Preprint at 10.1101/2024.10.02.616246 (2024).

19. Ranzoni, A. M. et al. Integrative Single-Cell RNA-Seq and ATAC-Seq Analysis of Human Developmental Hematopoiesis. Cell Stem Cell 28, 472–487.e7 (2021).

20. Jiang, P. et al. Single-cell ATAC-seq maps the comprehensive and dynamic chromatin accessibility landscape of CAR-T cell dysfunction. Leukemia 36, 2656–2668 (2022).

21. Janssens, J. et al. Decoding gene regulation in the fly brain. Nature 601, 630–636 (2022).

22. Johansen, N. J. et al. Evaluating Methods for the Prediction of Cell Type-Specific Enhancers in the Mammalian Cortex. 2024.08.21.609075 Preprint at 10.1101/2024.08.21.609075v3 (2025).

23. Ben-Simon, Y. et al. A suite of enhancer AAVs and transgenic mouse lines for genetic access to cortical cell types. 2024.06.10.597244 Preprint at 10.1101/2024.06.10.597244 (2024).

24. Cusanovich, D. A. et al. The cis-regulatory dynamics of embryonic development at single cell resolution. Nature 555, 538–542 (2018).

25. Hingerl, J. C. et al. scooby: Modeling multi-modal genomic profiles from DNA sequence at single-cell resolution. 2024.09.19.613754 Preprint at 10.1101/2024.09.19.613754 (2024).

26. Lal, A. et al. Decoding sequence determinants of gene expression in diverse cellular and disease states. 2024.10.09.617507 Preprint at 10.1101/2024.10.09.617507 (2025).

27. Hu, Y. et al. Multiscale footprints reveal the organization of cis-regulatory elements. Nature 1–8 (2025) doi:10.1038/s41586-024-08443-4.

28. Kaplow, I. M. et al. Relating enhancer genetic variation across mammals to complex phenotypes using machine learning. Science 380, eabm7993 (2023).

29. Hecker, N. et al. Enhancer-driven cell type comparison reveals similarities between the mammalian and bird pallium. Science 387, eadp3957 (2025).

30. Bravo González-Blas, C., et al. Single-cell spatial multi-omics and deep learning dissect enhancer-driven gene regulatory networks in liver zonation. Nat. Cell Biol. 26, 153–167 (2024).

31. Taskiran, I. I. et al. Cell-type-directed design of synthetic enhancers. Nature 626, 212–220 (2024).

32. Gosai, S. J. et al. Machine-guided design of cell-type-targeting cis-regulatory elements. Nature 634, 1211–1220 (2024).

33. de Almeida, B. P. et al. Targeted design of synthetic enhancers for selected tissues in the Drosophila embryo. Nature 626, 207–211 (2024).

34. Klie, A. et al. Predictive analyses of regulatory sequences with EUGENe. Nat. Comput. Sci. 3, 946–956 (2023).

35. Lal, A., Gunsalus, L., Nair, S., Biancalani, T. & Eraslan, G. gReLU: A comprehensive framework for DNA sequence modeling and design. 2024.09.18.613778 Preprint at 10.1101/2024.09.18.613778 (2024).

36. Chen, K. M., Cofer, E. M., Zhou, J. & Troyanskaya, O. G. Selene: a PyTorch-based deep learning library for sequence data. Nat. Methods 16, 315–318 (2019).

37. Virshup, I. et al. The scverse project provides a computational ecosystem for single-cell omics data analysis. Nat. Biotechnol. 41, 604–606 (2023).

38. Granja, J. M. et al. ArchR is a scalable software package for integrative single-cell chromatin accessibility analysis. Nat. Genet. 53, 403–411 (2021).

39. Bravo González-Blas, C., et al. SCENIC+: single-cell multiomic inference of enhancers and gene regulatory networks. Nat. Methods 20, 1355–1367 (2023).

40. Zhang, K., Zemke, N. R., Armand, E. J. & Ren, B. A fast, scalable and versatile tool for analysis of single-cell omics data. Nat. Methods 21, 217–227 (2024).

41. Bravo González-Blas, C., et al. cisTopic: cis-regulatory topic modeling on single-cell ATAC-seq data. Nat. Methods 16, 397–400 (2019).

42. Pampari, A., et al. Bias factorized, base-resolution deep learning models of chromatin accessibility reveal cis-regulatory sequence syntax, transcription factor footprints and regulatory variants. Zenodo 10.5281/zenodo.10396047 (2023).

43. p-koo/tfomics. https://github.com/p-koo/tfomics/tree/master?tab=readme-ov-file.

44. GitHub - jmschrei/tfmodisco-lite: A lite implementation of tfmodisco, a motif discovery algorithm for genomics experiments. https://github.com/jmschrei/tfmodisco-lite.

45. Shrikumar, A., et al. Technical Note on Transcription Factor Motif Discovery from Importance Scores (TF-MoDISco) version 0.5.6.5. Preprint at 10.48550/arXiv.1811.00416 (2020).

46. jmschrei/tangermeme: Biological sequence analysis for the modern age. https://github.com/jmschrei/tangermeme/tree/main.

47. Zemke, N. R. et al. Conserved and divergent gene regulatory programs of the mammalian neocortex. Nature 624, 390–402 (2023).

48. Hounkpe, B. W., Chenou, F., de Lima, F. & De Paula, E. V. HRT Atlas v1.0 database: redefining human and mouse housekeeping genes and candidate reference transcripts by mining massive RNA-seq datasets. Nucleic Acids Res. 49, D947–D955 (2021).

49. De Rop, F. V. et al. Systematic benchmarking of single-cell ATAC-sequencing protocols. Nat. Biotechnol. 42, 916–926 (2024).

50. Nutt, S. L. & Kee, B. L. The Transcriptional Regulation of B Cell Lineage Commitment. Immunity 26, 715–725 (2007).

51. Spicuglia, S. et al. TCRα enhancer activation occurs via a conformational change of a pre-assembled nucleo-protein complex. EMBO J. 19, 2034–2045 (2000).

52. Panne, D., Maniatis, T. & Harrison, S. C. An Atomic Model of the Interferon-β Enhanceosome. Cell 129, 1111–1123 (2007).

53. Thanos, D. & Maniatis, T. Virus induction of human IFNβ gene expression requires the assembly of an enhanceosome. Cell 83, 1091–1100 (1995).

54. Garrett-Sinha, L. A. Review of Ets1 structure, function, and roles in immunity. Cell. Mol. Life Sci. 70, 3375–3390 (2013).

55. Schmidl, C. et al. The enhancer and promoter landscape of human regulatory and conventional T-cell subpopulations. Blood 123, e68–78 (2014).

56. Saint-André, V. et al. Models of human core transcriptional regulatory circuitries. Genome Res. 26, 385–396 (2016).

57. Heinz, S. et al. Transcription Elongation Can Affect Genome 3D Structure. Cell 174, 1522–1536.e22 (2018).

58. Trompouki, E. et al. Lineage regulators direct BMP and Wnt pathways to cell-specific programs during differentiation and regeneration. Cell 147, 577–589 (2011).

59. Zhang, J. et al. An integrative ENCODE resource for cancer genomics. Nat. Commun. 11, 3696 (2020).

60. Hollenhorst, P. C. et al. DNA specificity determinants associate with distinct transcription factor functions. PLoS Genet. 5, e1000778 (2009).

61. Valouev, A. et al. Genome-wide analysis of transcription factor binding sites based on ChIP-Seq data. Nat. Methods 5, 829–834 (2008).

62. Gheorghe, M. et al. A map of direct TF–DNA interactions in the human genome. Nucleic Acids Res. 47, e21 (2019).

63. Fedl, A. S. et al. Transcriptional function of E2A, Ebf1, Pax5, Ikaros and Aiolos analyzed by in vivo acute protein degradation in early B cell development. Nat. Immunol. 25, 1663–1677 (2024).

64. Dawson, M. A. The cancer epigenome: Concepts, challenges, and therapeutic opportunities. Science 355, 1147–1152 (2017).

65. Tirosh, I. & Suva, M. L. Cancer cell states: Lessons from ten years of single-cell RNA-sequencing of human tumors. Cancer Cell 42, 1497–1506 (2024).

66. Barkley, D. et al. Cancer cell states recur across tumor types and form specific interactions with the tumor microenvironment. Nat. Genet. 54, 1192–1201 (2022).

67. Kinker, G. S. et al. Pan-cancer single-cell RNA-seq identifies recurring programs of cellular heterogeneity. Nat. Genet. 52, 1208–1218 (2020).

68. Terekhanova, N. V. et al. Epigenetic regulation during cancer transitions across 11 tumour types. Nature 623, 432–441 (2023).

69. Mauduit, D. et al. Analysis of long and short enhancers in melanoma cell states. eLife 10, e71735 (2021).

70. Dunham, I. et al. An integrated encyclopedia of DNA elements in the human genome. Nature 489, 57–74 (2012).

71. Schnöller, L. E. et al. Systematic in vitro analysis of therapy resistance in glioblastoma cell lines by integration of clonogenic survival data with multi-level molecular data. Radiat. Oncol. 18, 51 (2023).

72. Goudarzi, K. M. et al. Reduced Expression of PROX1 Transitions Glioblastoma Cells into a Mesenchymal Gene Expression Subtype. Cancer Res. 78, 5901–5916 (2018).

73. Kircher, M. et al. Saturation mutagenesis of twenty disease-associated regulatory elements at single base-pair resolution. Nat. Commun. 10, 3583 (2019).

74. Verfaillie, A. et al. Decoding the regulatory landscape of melanoma reveals TEADS as regulators of the invasive cell state. Nat. Commun. 6, 6683 (2015).

75. Wouters, J. et al. Robust gene expression programs underlie recurrent cell states and phenotype switching in melanoma. Nat. Cell Biol. 22, 986–998 (2020).

76. Wang, L. et al. The Phenotypes of Proliferating Glioblastoma Cells Reside on a Single Axis of Variation. Cancer Discov. 9, 1708–1719 (2019).

77. Schwessinger, R., Deasy, J., Woodruff, R. T., Young, S. & Branson, K. M. Single-cell gene expression prediction from DNA sequence at large contexts. 2023.07.26.550634 Preprint at 10.1101/2023.07.26.550634 (2023).

78. Murphy, A. E., Beardall, W., Rei, M., Phuycharoen, M. & Skene, N. G. Predicting cell type-specific epigenomic profiles accounting for distal genetic effects. Nat. Commun. 15, 9951 (2024).

79. Li, Y. E. et al. An atlas of gene regulatory elements in adult mouse cerebrum. Nature 598, 129–136 (2021).

80. Sun, K. et al. Mapping the chromatin accessibility landscape of zebrafish embryogenesis at single-cell resolution by SPATAC-seq. Nat. Cell Biol. 26, 1187–1199 (2024).

81. Begeman, I. J., Emery, B., Kurth, A. & Kang, J. Regeneration and developmental enhancers are differentially compatible with minimal promoters. Dev. Biol. 492, 47–58 (2022).

82. Wasserman, W. W. & Sandelin, A. Applied bioinformatics for the identification of regulatory elements. Nat. Rev. Genet. 5, 276–287 (2004).

83. Arnosti, D. N. & Kulkarni, M. M. Transcriptional enhancers: Intelligent enhanceosomes or flexible billboards? J. Cell. Biochem. 94, 890–898 (2005).

84. Long, H. K., Prescott, S. L. & Wysocka, J. Ever-Changing Landscapes: Transcriptional Enhancers in Development and Evolution. Cell 167, 1170–1187 (2016).

85. Matharu, N. & Ahituv, N. Modulating gene regulation to treat genetic disorders. Nat. Rev. Drug Discov. 19, 757–775 (2020).

86. Chandra, N. A., Hu, Y., Buenrostro, J. D., Mostafavi, S. & Sasse, A. Refining the cis-regulatory grammar learned by sequence-to-activity models by increasing model resolution. bioRxiv 2025–01 (2025).

87. Huey, J. D. & Abdennur, N. Bigtools: a high-performance BigWig and BigBed library in Rust. Bioinformatics 40, btae350 (2024).

88. Virshup, I., Rybakov, S., Theis, F. J., Angerer, P. & Wolf, F. A. anndata: Access and store annotated data matrices. J. Open Source Softw. 9, 4371 (2024).

89. Sundararajan, M., Taly, A. & Yan, Q. Axiomatic Attribution for Deep Networks. in Proceedings of the 34th International Conference on Machine Learning 3319–3328 (PMLR, 2017).

90. Majdandzic, A., Rajesh, C. & Koo, P. K. Correcting gradient-based interpretations of deep neural networks for genomics. Genome Biol. 24, 109 (2023).

91. Erion, G., Janizek, J. D., Sturmfels, P., Lundberg, S. M. & Lee, S.-I. Improving performance of deep learning models with axiomatic attribution priors and expected gradients. Nat. Mach. Intell. 3, 620–631 (2021).

92. Korsunsky, I. et al. Fast, sensitive and accurate integration of single-cell data with Harmony. Nat. Methods 16, 1289–1296 (2019).

93. Langmead, B. & Salzberg, S. L. Fast gapped-read alignment with Bowtie 2. Nat. Methods 9, 357–359 (2012).

94. Zhang, Y. et al. Model-based Analysis of ChIP-Seq (MACS). Genome Biol. 9, R137 (2008).

95. Ramírez, F. et al. deepTools2: a next generation web server for deep-sequencing data analysis. Nucleic Acids Res. 44, W160–W165 (2016).

96. Hammal, F., de Langen, P., Bergon, A., Lopez, F. & Ballester, B. ReMap 2022: a database of Human, Mouse, Drosophila and Arabidopsis regulatory regions from an integrative analysis of DNA-binding sequencing experiments. Nucleic Acids Res. 50, D316–D325 (2022).

