## Supplementary Information for "CREsted: modeling genomic and synthetic cell type-specific enhancers across tissues and species"

Niklas Kempynck[1,2,3,7], Seppe De Winter [1,2,3,7], Casper H. Blaauw[1,2,3,4], Vasileios Konstantakos[1,2,3], Sam Dieltiens[1,2,3], Eren Can Ekşi[1,2,3], Valérie Bercier[2,5], Ibrahim I. Taskiran[1,2,3,$], Gert Hulselmans[1,2,3,6], Katina Spanier[1,2,3], Valerie Christiaens[1,2,3,6], Ludo Van Den Bosch[2,5], Lukas Mahieu[1,2,3,6], Stein Aerts[1,2,3,6,*]

This file contains:

- Supplementary Figures S1-16
- Supplementary Tables S1-2

### Supplementary Figures

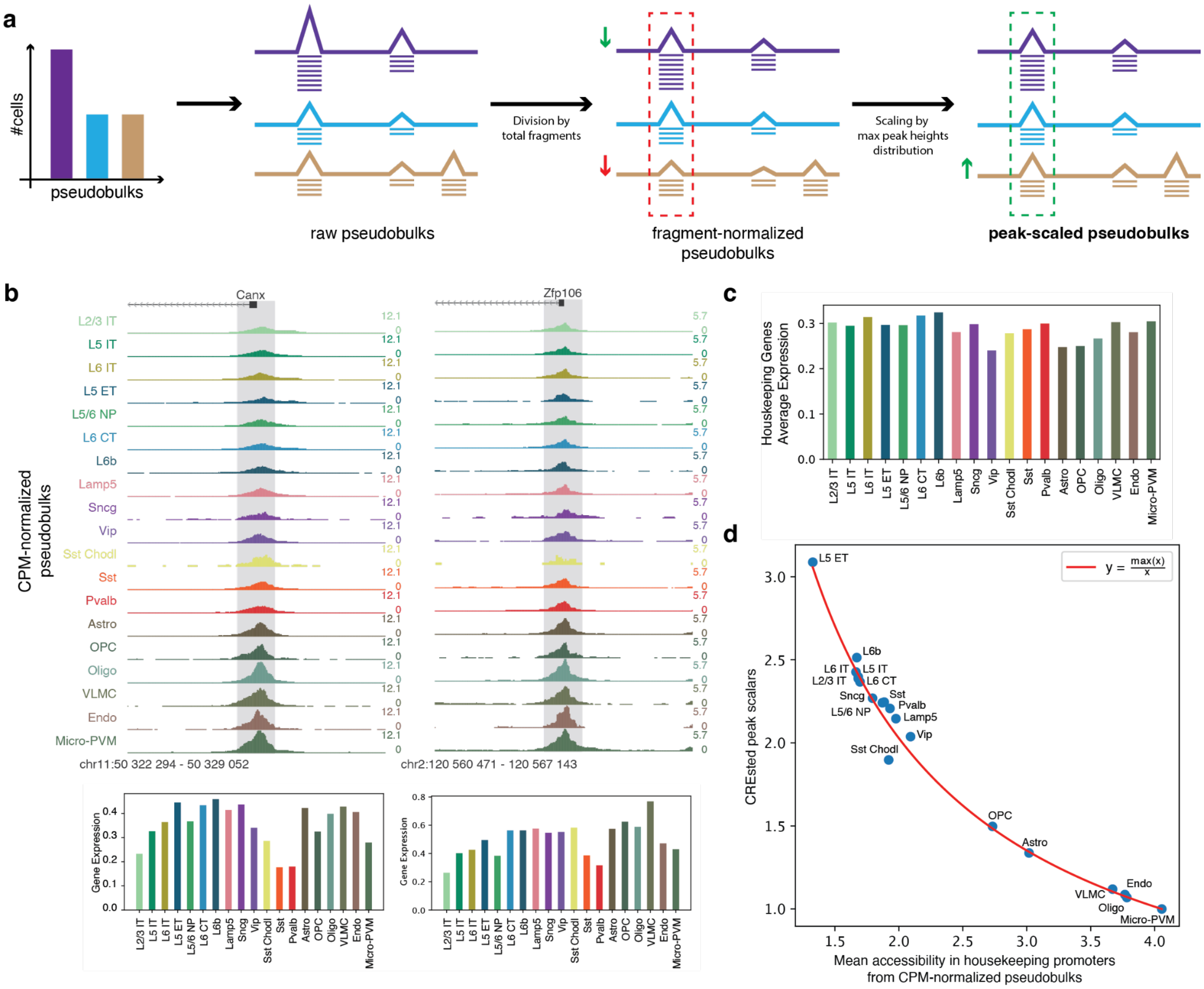


#### Figure S1. Overview of peak scaling method.

(**a**) Overview of the peak normalization method. (**b**) Peak heights over all cell types of the housekeeping *Canx* and *Zfp106* genes from standard CPM-normalized scATAC-seq tracks (top). Normalized expression of both genes over all cell types (bottom). (**c**) Average expression for all housekeeping genes over all cell types. (**d**) Comparison of average peak heights in housekeeping promoters and the peak scalars used to counterbalance them. The expected curve is shown in red.

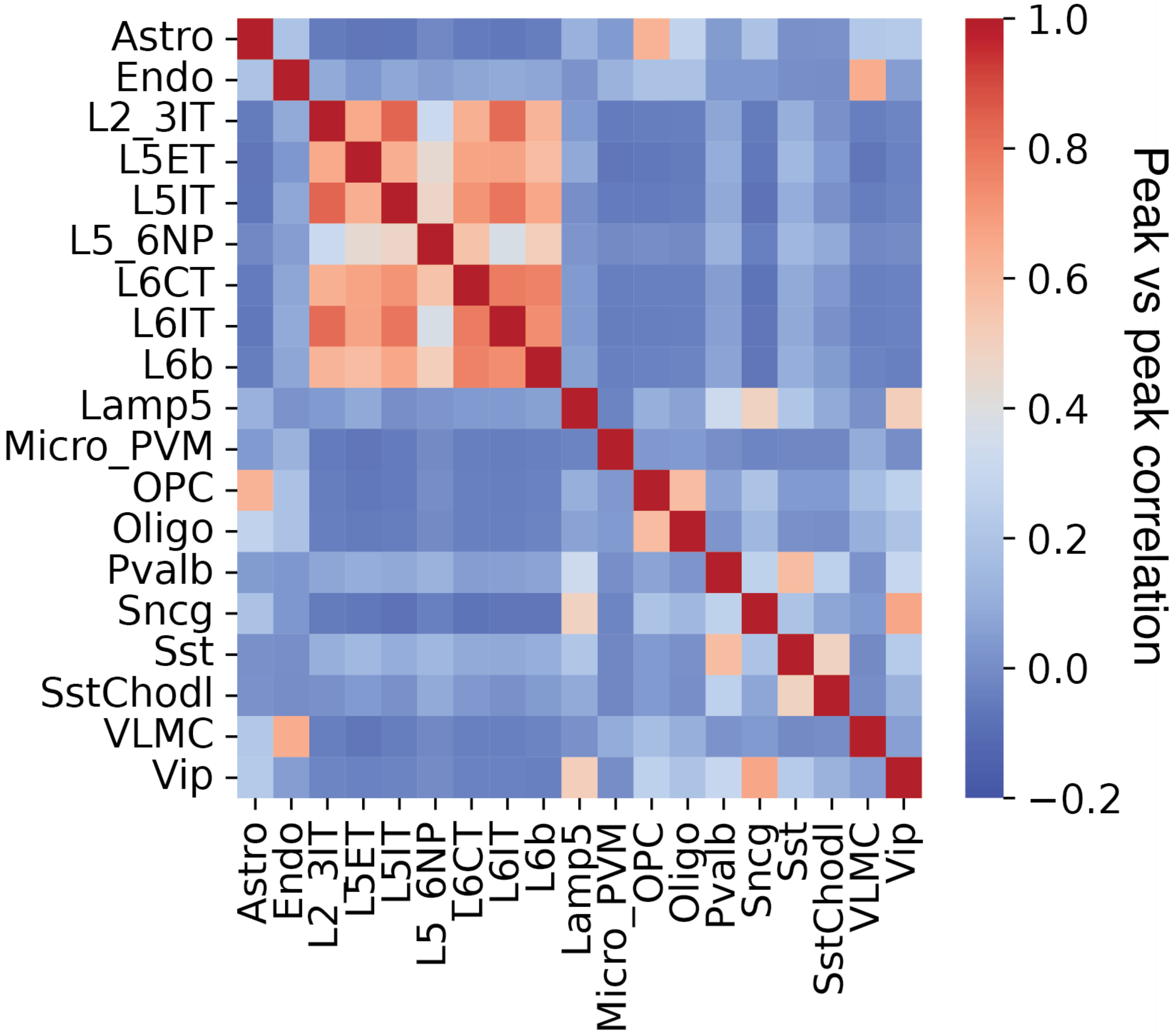


Figure S2. Correlation of peak heights for mouse motor cortex cell types.

Heatmap indicating correlation of peak heights across cell types from a set of cell type-specific regions.

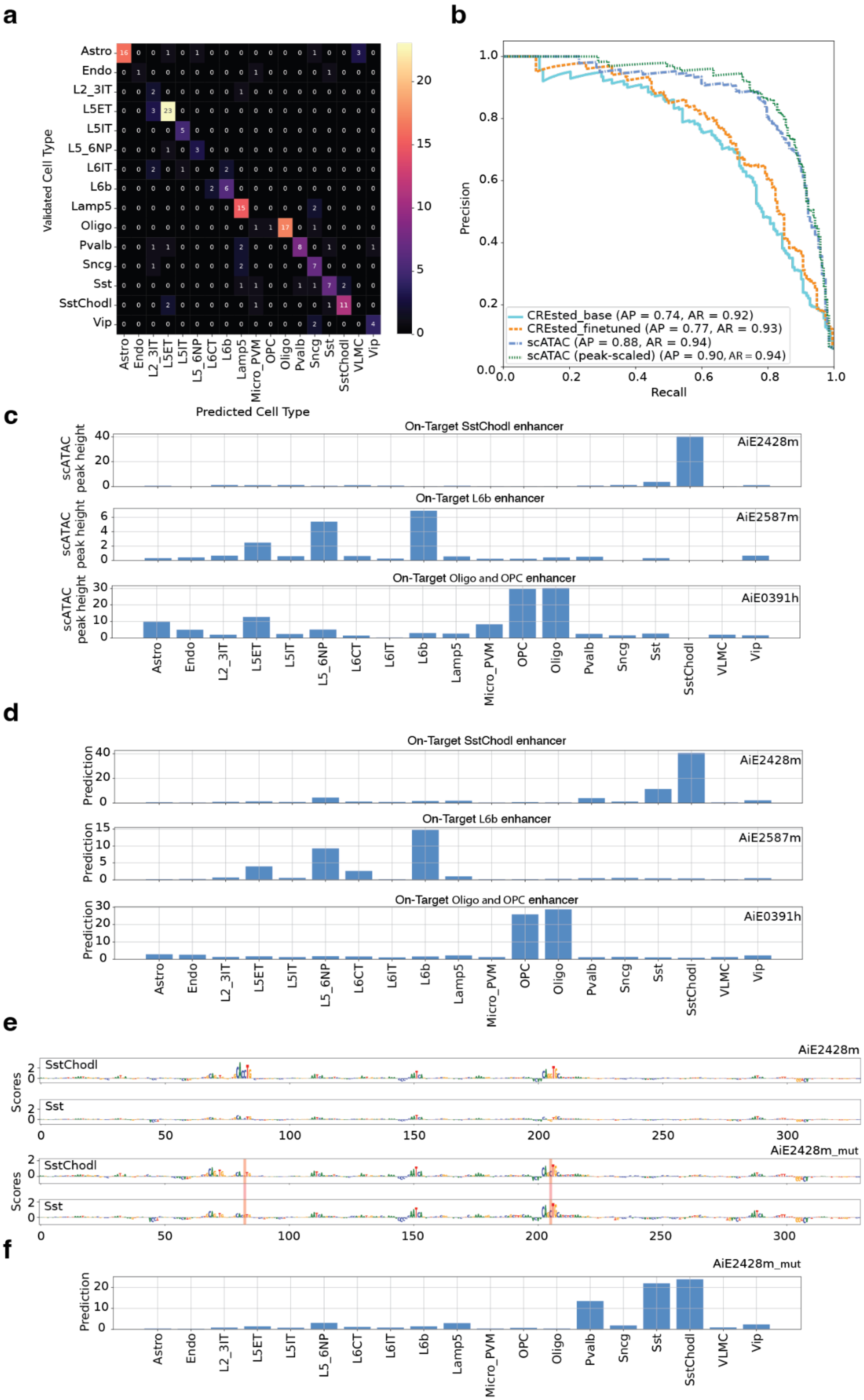
a
b
Validated Cell Type
Predicted Cell Type
Astro
Endo
L2_3IT
L5ET
L5IT
L5_6NP
L6IT
L6b
Lamp5
Oligo
Pvalb
Sncg
Sst
SstChodl
Vip
L6CT
Micro_PVM
OPC
VLMC
Precision
Recall
CREsted_base (AP = 0.74, AR = 0.92)
CREsted_finetuned (AP = 0.77, AR = 0.93)
scATAC (AP = 0.88, AR = 0.94)
scATAC (peak-scaled) (AP = 0.90, AR = 0.94)
c
On-Target SstChodl enhancer
AiE2428m
scATAC peak height
On-Target L6b enhancer
AiE2587m
On-Target Oligo and OPC enhancer
AiE0391h
d
Prediction
e
Scores
SstChodl
Sst
AiE2428m_mut
f

#### Figure S3. CREsted on *in vivo* validated mouse cortex enhancers.

(**a**) Heatmap of predicted labels from the maximum prediction per region and the targeted validated cell types for a set of *in vivo* validated enhancers. (**b**) Averaged multi-label precision-recall curve for predicting the set of validated enhancers indicating specificity per cell types. Specificity was calculated for scATAC peaks (scaled and non-scaled), and for CREsted predictions (basemodel and fine-tuned). (**c**) scATAC peak heights for three example enhancer regions. (**d**) Predictions for three example enhancer regions. (**e**) Contribution scores for the AiE2428m SstChodl enhancer, before (top) and after (bottom) E-box motif mutation. (**f**) Prediction scores for mutated AiE2428m enhancer.

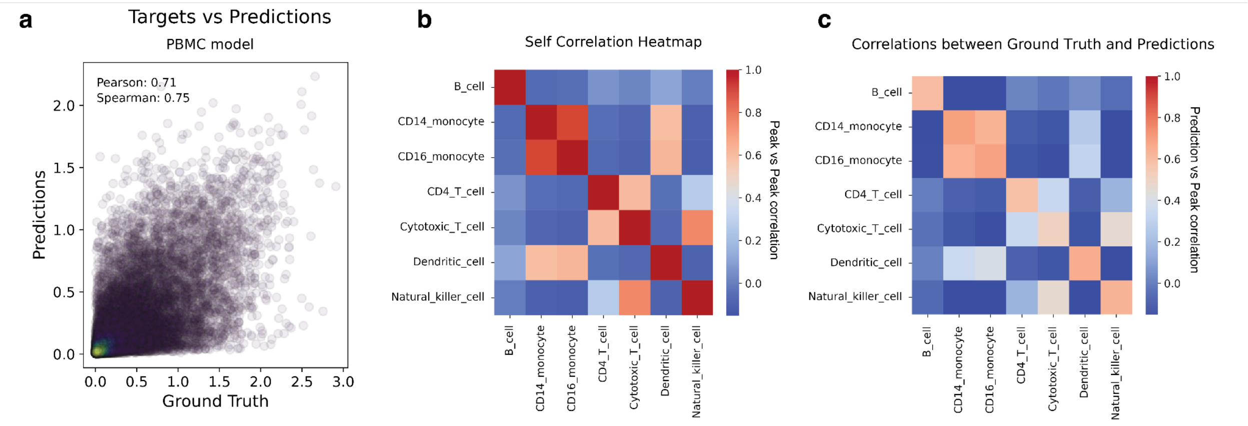


#### Figure S4. Fine-tuned CREsted PBMC model performance overview.

(**a**) Scatter plot of all predicted targets and predictions for all specific test set regions over all classes. (**b**) Accessibility correlation of specific test regions across cell types. (**c**) Correlation of predictions and accessibility over cell types for specific test set regions.

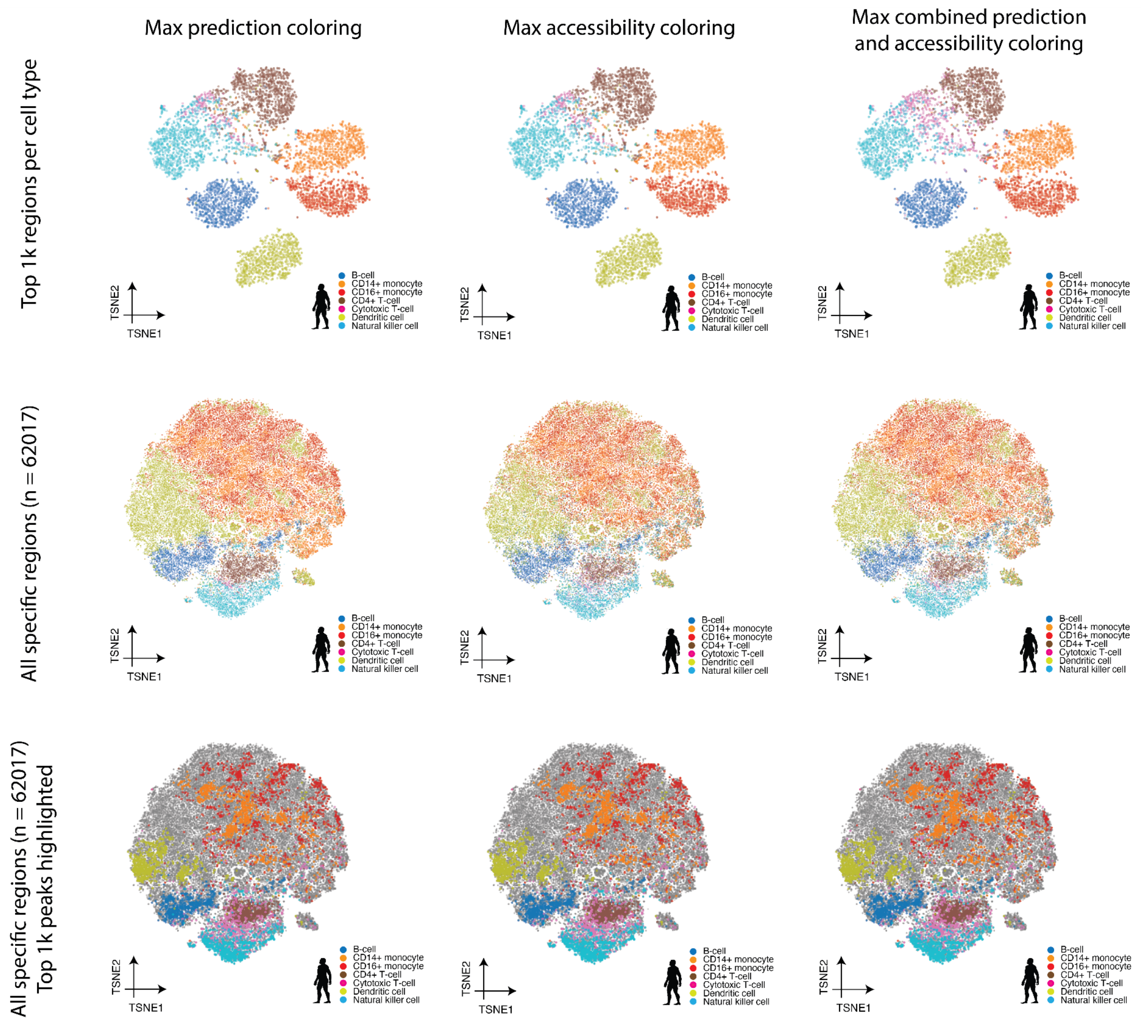


#### Figure S5. Region embeddings from DeepPBMC.

Embeddings of the second-to-last DeepPBMC model layer of the top 1,000 regions per cell type (top), of all cell type-specific peaks (middle row) and of all cell type-specific peaks with only the top 1,000 regions per cell type highlighted (bottom). The left column colors regions by their max DeepPBMC prediction value, the middle column by their max peak height, and the right column by their average.

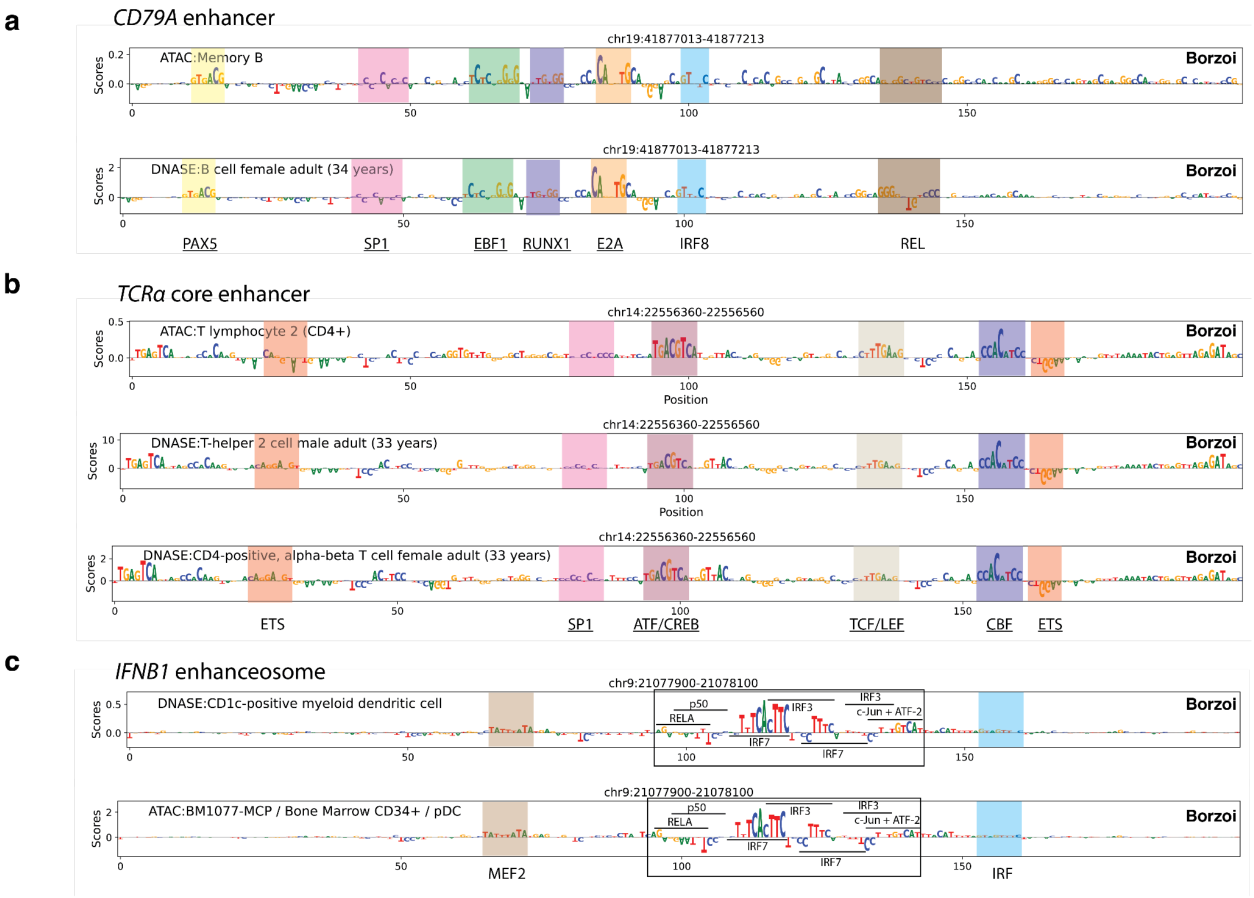


### Figure S6. Base Borzoi contribution scores on validated enhancers.

Contribution scores from the base Borzoi model for the *CD79A* (hg38 chr19:41,876,056-41,878,170) (**a**) and *TCRα* core (hg38 chr14:22,555,403-22,557,517) (**b**) enhancers and the *IFNB1* enhanceosome (hg38 chr9:21,076,963-21,079,077) (**c**) for a selection of relevant classes. Identified motifs are highlighted and annotated. Validated TFBS are underlined.

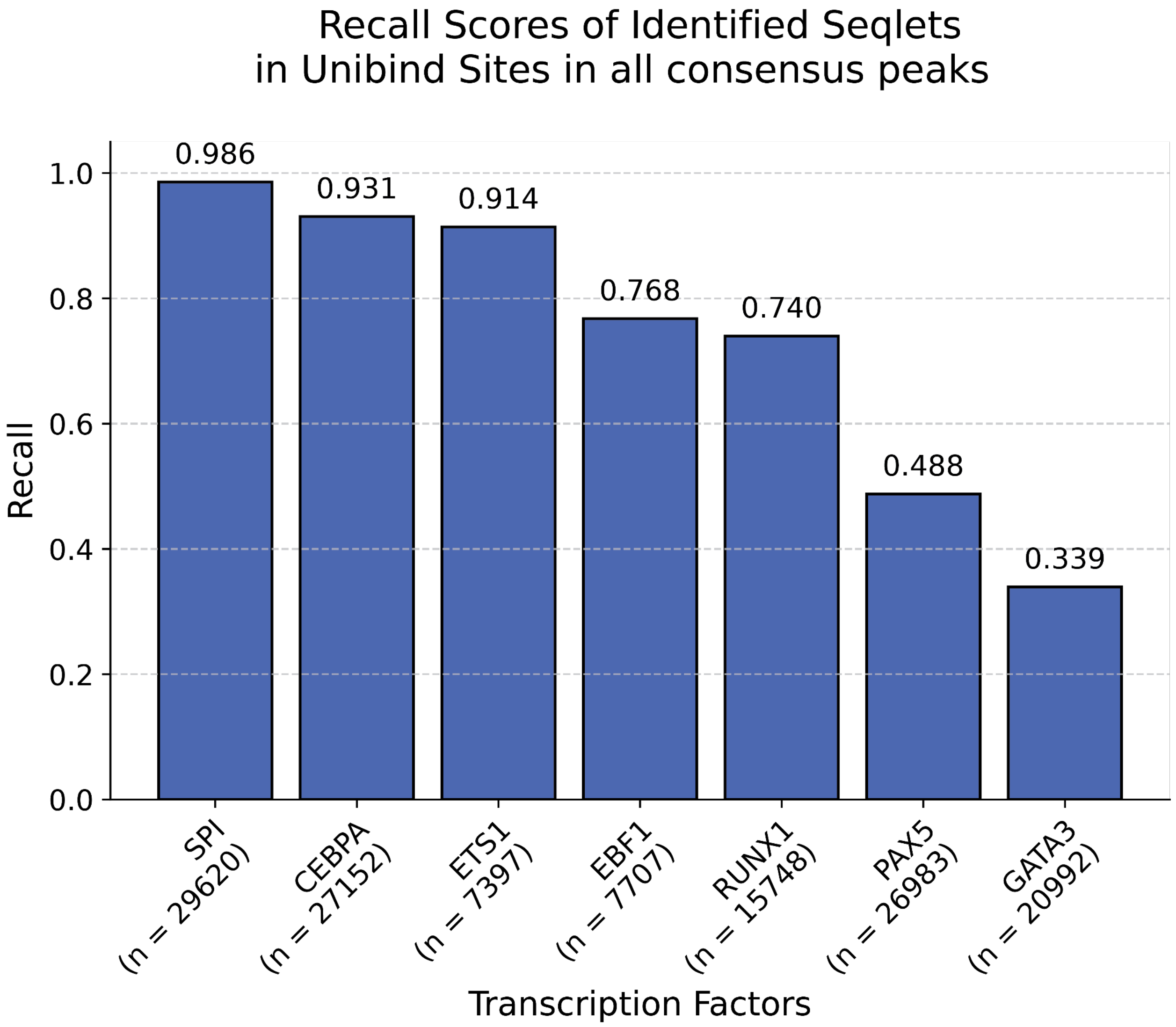


Figure S7. Recall of UniBind sites identified by DeepPBMC in all consensus peaks.

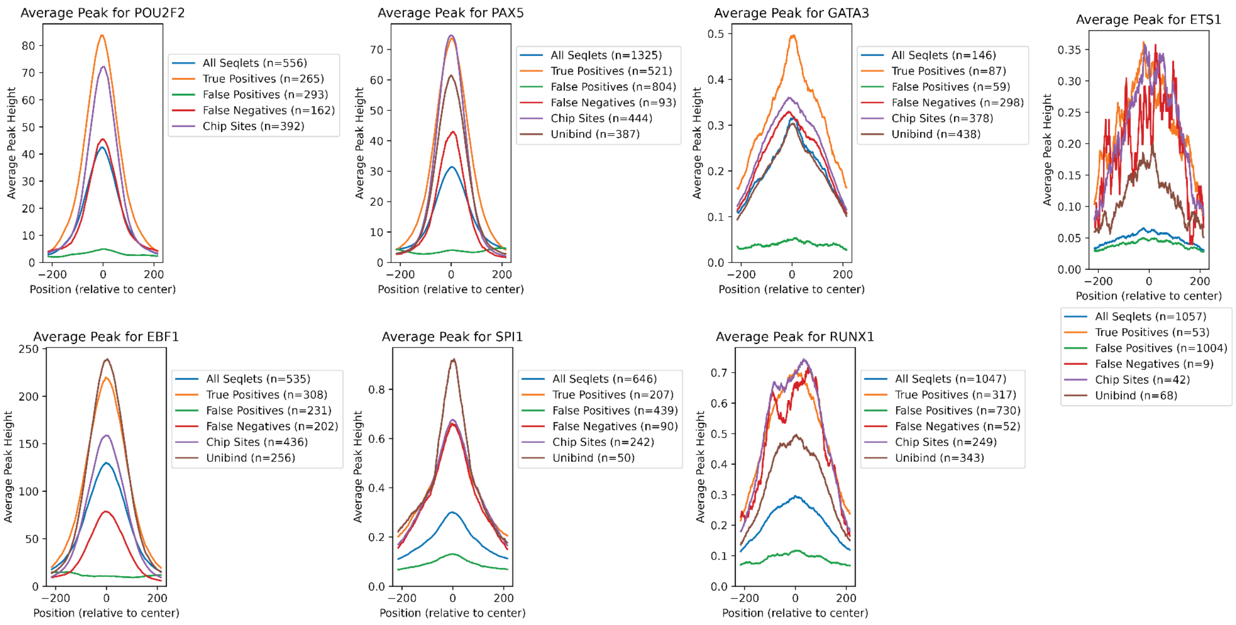


#### Figure S8. ChIP-seq peak heights for a selection of TFs

The aggregated peak heights of all identified seqlets, true positive seqlets, false positive seqlets, false negative ChIP peaks, and all ChIP-seq and UniBind hits in the top 1,000 regions per cell type are shown per TF.

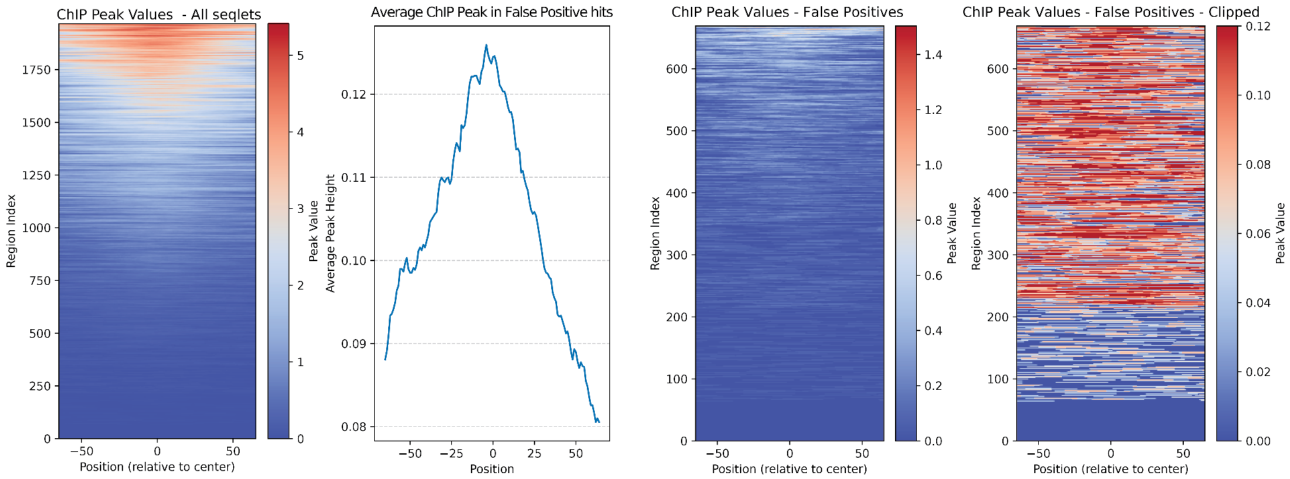


#### Figure S9. ChIP-seq peak heights for seqlet instances

Carrot plot of ChIP-seq peak heights of all identified CEBPA seqlets (left), the aggregated ChIP-seq peak for all false positive CEBPA seqlets (second to left), carrot plot of ChIP-seq peak heights of false positive CEBPA seqlets (second to right) and a clipped carrot plot of ChIP-seq peak heights of false positive CEBPA seqlets.

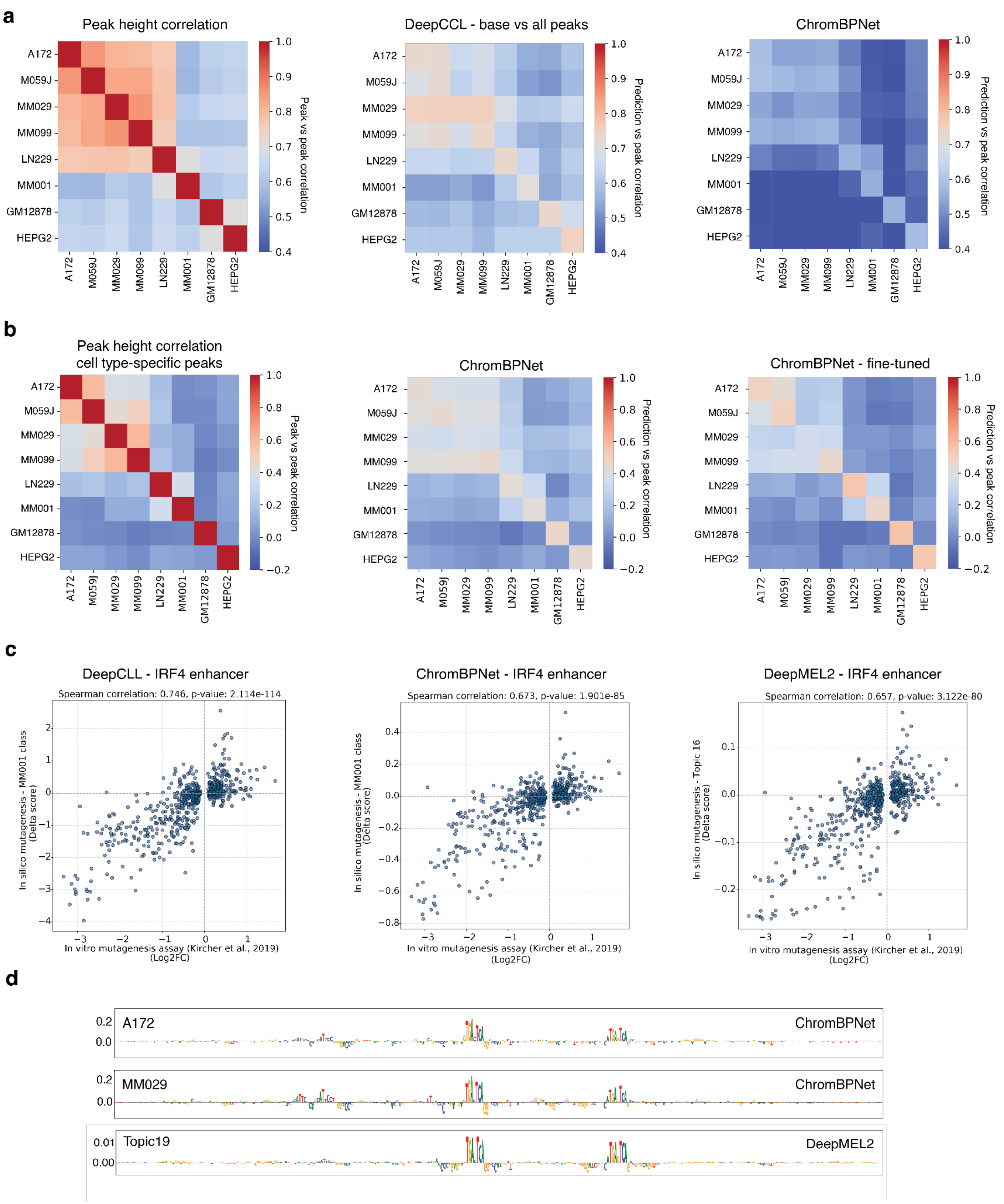

a
Peak height correlation
DeepCCL - base vs all peaks
ChromBPNet
A172
M059J
MM029
MM099
LN229
MM001
GM12878
HEPG2
Peak vs peak correlation
Prediction vs peak correlation
b
Peak height correlation
cell type-specific peaks
ChromBPNet
ChromBPNet - fine-tuned
c
DeepCLL - IRF4 enhancer
Spearman correlation: 0.746, p-value: 2.114e-114
In silico mutagenesis - MM001 class
(Delta score)
In vitro mutagenesis assay (Kircher et al., 2019)
(Log2FC)
ChromBPNet - IRF4 enhancer
Spearman correlation: 0.673, p-value: 1.901e-85
DeepMEL2 - IRF4 enhancer
Spearman correlation: 0.657, p-value: 3.122e-80
In silico mutagenesis - Topic 16
(Delta score)
d
A172
MM029
Topic19
ChromBPNet
DeepMEL2

#### Figure S10. DeepCCL, ChromBPNet and DeepMEL2 performance comparison.

(**a**) For the set of all test set regions, the log-transformed peak height Pearson correlation across cell types (left), the Pearson correlation between the log-transformed DeepCLL base model predictions and peak values (middle) and the Pearson correlation between the combined ChromBPNet model predictions and peak values (right). (**b**) For the set of cell type-specific test set regions, the log-transformed peak height Pearson correlation across cell types (left), the Pearson correlation between the log-transformed combined ChromBPNet model predictions and peak values (middle) and the Pearson correlation between the combined ChromBPNet model, fine-tuned on specific regions, predictions and peak values (right). (**c**) Scatter plot comparison between ISM and in vitro mutagenesis values[73] for the IRF4 enhancer in MEL-classes (MM001 in DeepCCL and ChromBPNet, and Topic 16 in DeepMEL2). (**d**) Contribution scores from ChromBPNet and DeepMEL2 for the identified intronic *AXL* region.

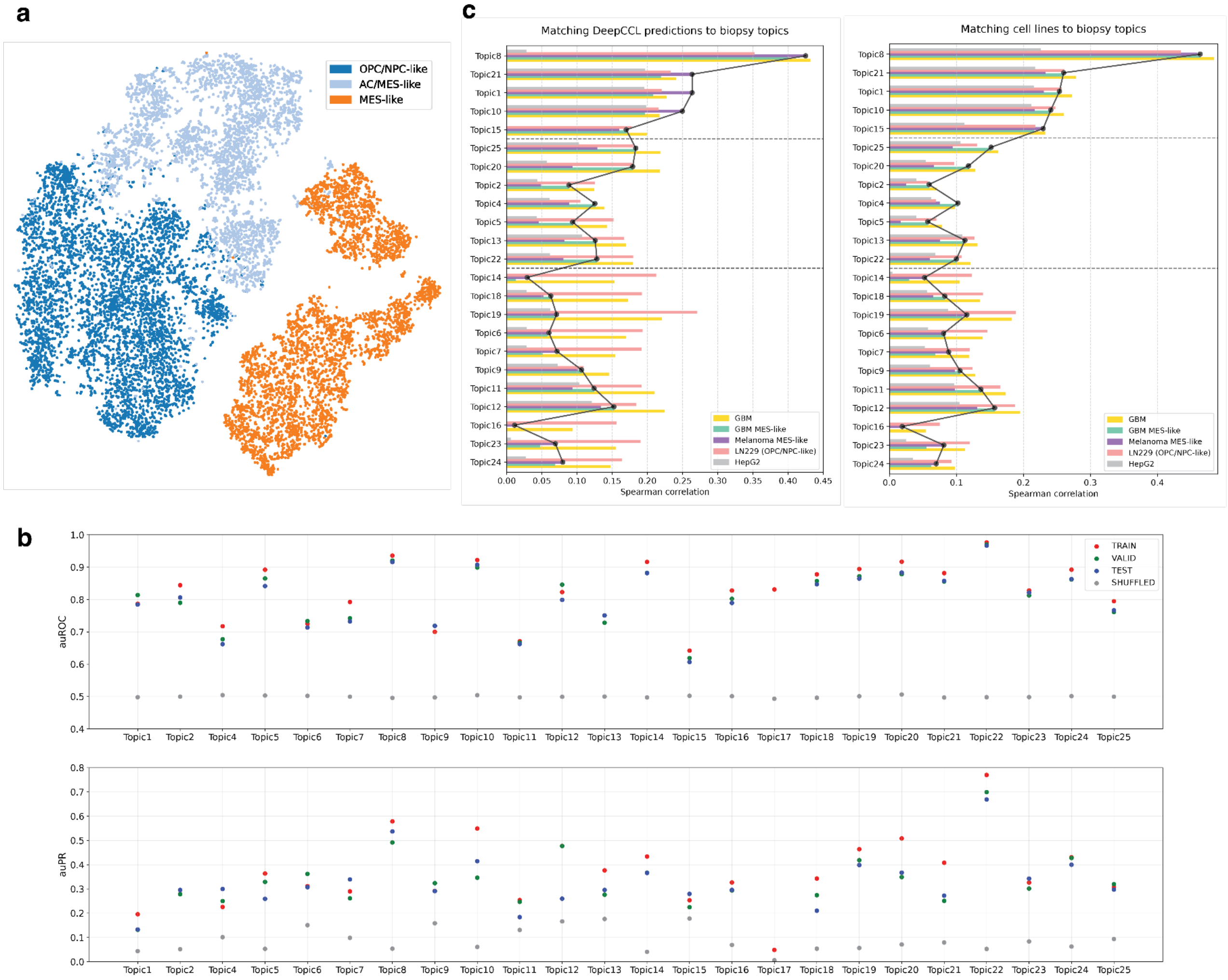


### Figure S11. Comparison between DeepCCL cell lines and DeepGlioma topics.

(**a**) Manually annotated clusters for the Wang *et al*. glioma biopsy data. (**b**) Area under receiver operating characteristic curve (auROC) (top) and area under PR-curve (bottom) per topic for the different splits for the DeepGlioma model. (**c**) Spearman correlation between DeepCCL predictions / cell line accessibility and pseudobulked Topic bigwig files obtained from the pycisTopic analysis. GBM: average prediction/accessibility of A172, M059J, and LN229. GBM MES-like: average prediction/accessibility of A172 and M059J. Melanoma MES-like: average prediction/accessibility of MM029 and MM099. Line plot illustrates the max MES-like correlation across the GBM MES-like and Melanoma MES-like bars.

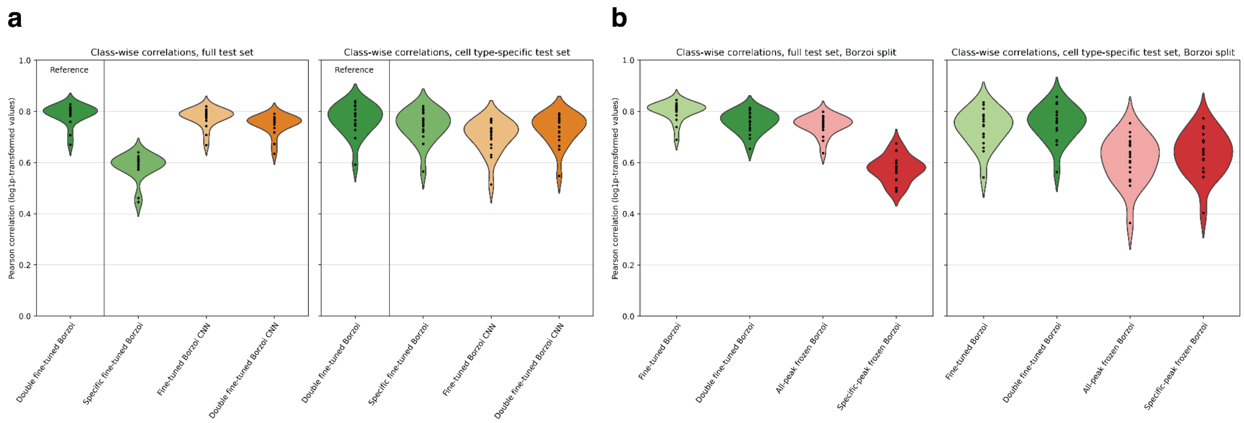


### Figure S12. Model performance of additional Borzoi transfer learned models

Violin plots showing Pearson correlation values between the log1p-transformed test sets (all consensus peaks and cell type-specific peaks respectively) and corresponding predictions, calculated across the peaks for each cell type. **(a)** Comparison of the best-performing transfer learned model (Double fine-tuned Borzoi) with other transfer learning approaches on the *r* per cell type (n=19). **(b)** Correlation values per cell type (n=19) for models trained and evaluated analogously to fig. 5b, but using a train/validation/test split based on the Borzoi folds instead.

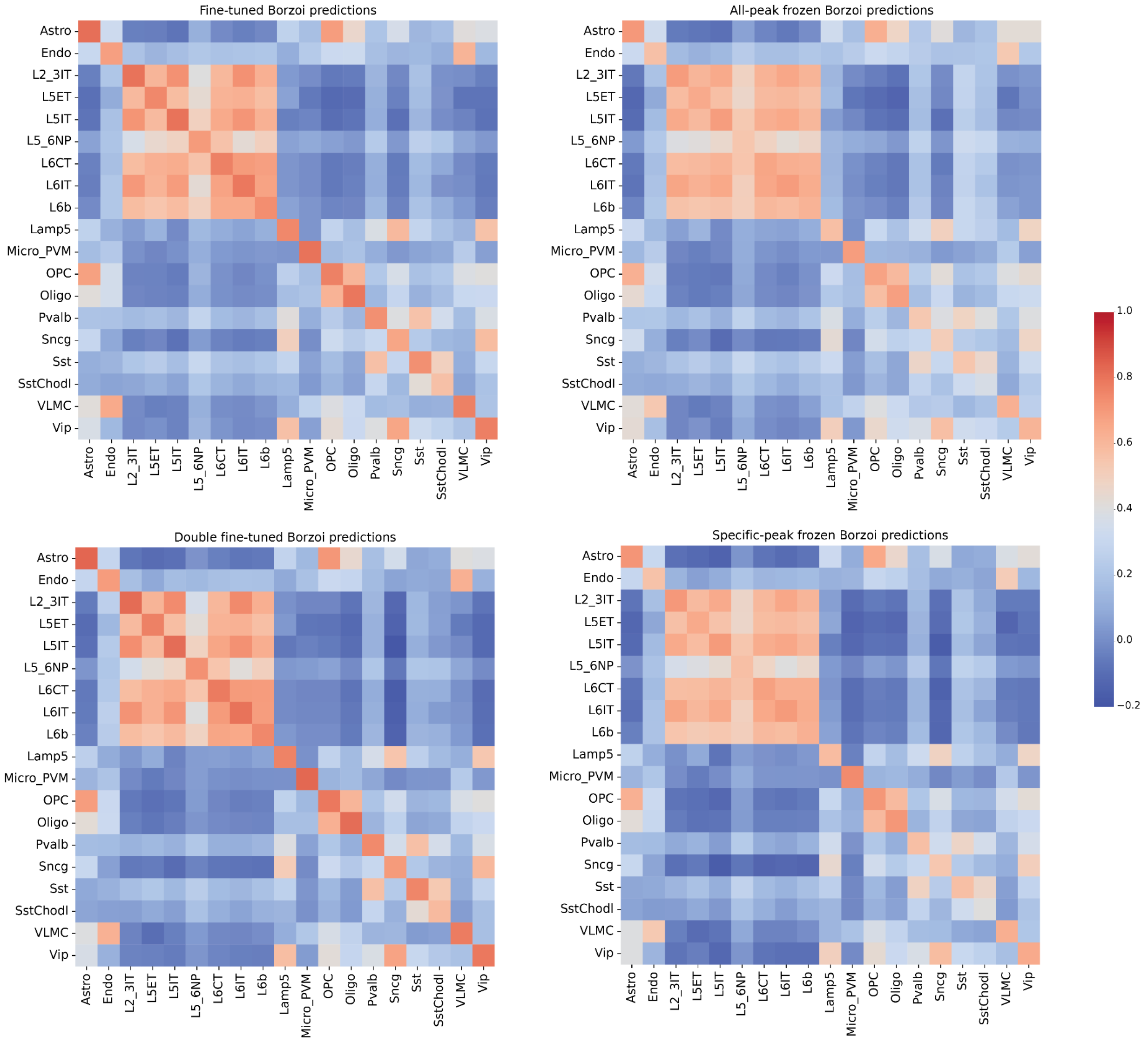


### Figure S13. Correlation heatmaps of Borzoi transfer learned models

Heatmaps of Pearson correlations for log1p-transformed cell type-specific test set peaks, analogous to Fig. 2c, for the predictions from the transfer learned models from Fig. 5b, comparing each cell type's peaks against each cell type's predictions.

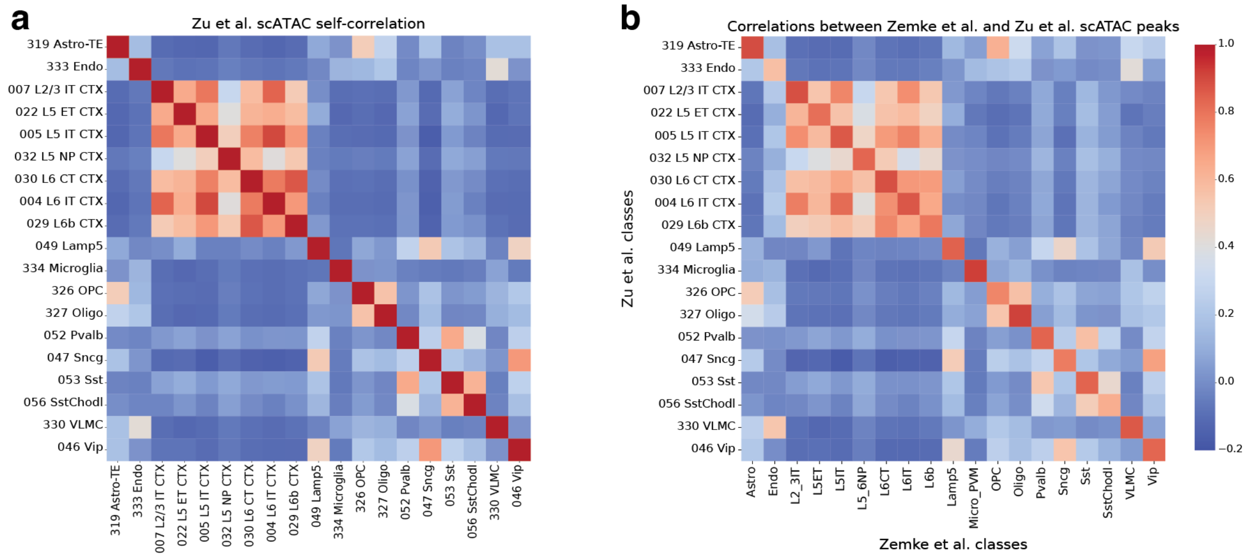


### Figure S14. Correlations of peak heights of the Zu et al. mouse brain scATAC dataset.

**(a)** Heatmap showing the self-correlation (Pearson correlations of log1p-transformed data) between selected Zu et al. pseudobulk scATAC classes, based on the full cell type-specific set. **(b)** Heatmap showing the correlations between Zu et al. and Zemke et al. scATAC pseudobulk classes, based on the full cell-type specific set.

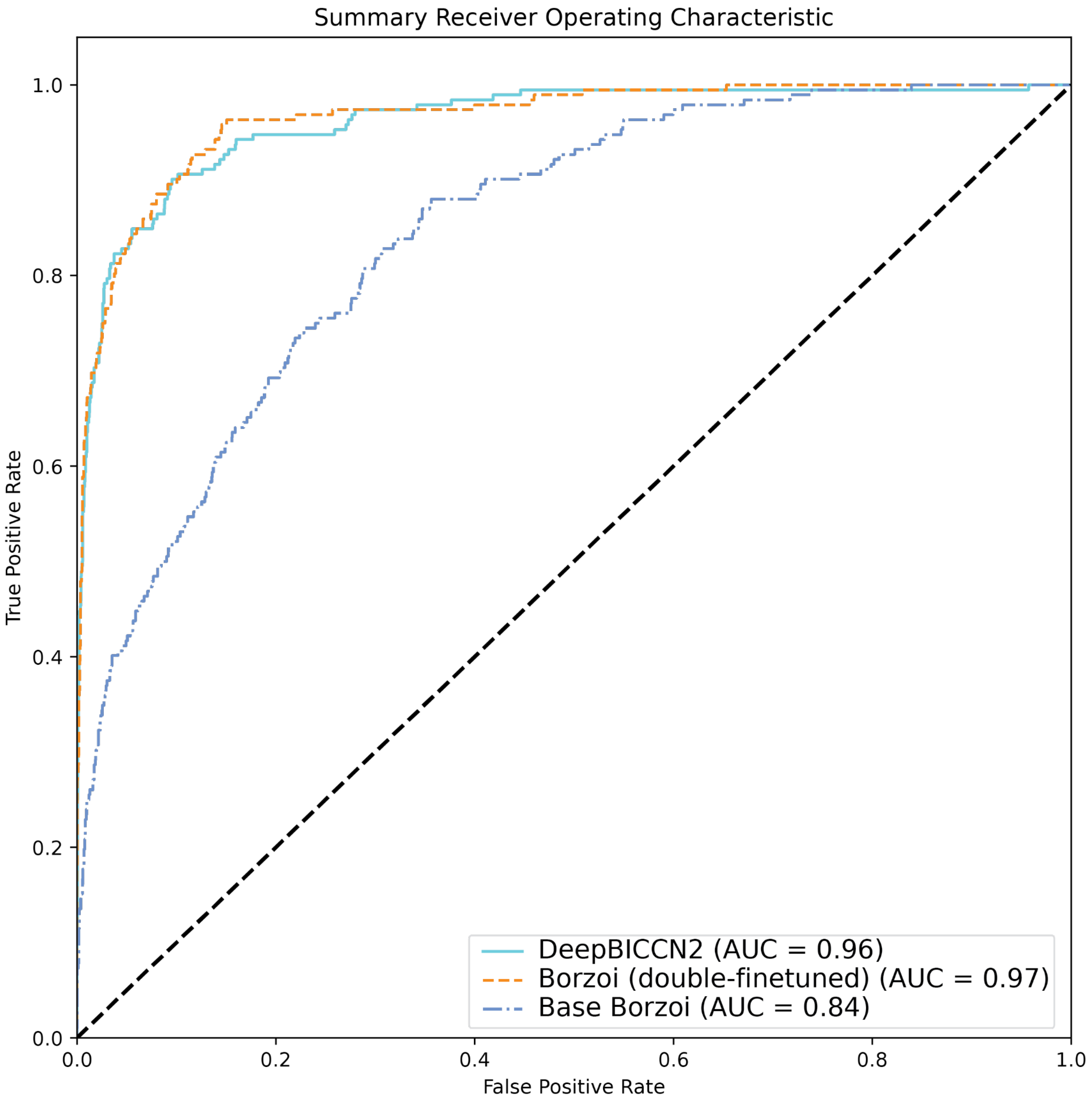


Figure S15. ROC curve of specificity predictions for validated enhancers.

Comparison of DeepBICCN2, base Borzoi, and the double fine-tuned Borzoi model on the Zemke *et al.* mouse motor cortex data. ROC curve was made on the true and false positive rate of the specificity of the predictions on a set of 171 *in vivo* validated enhancers.

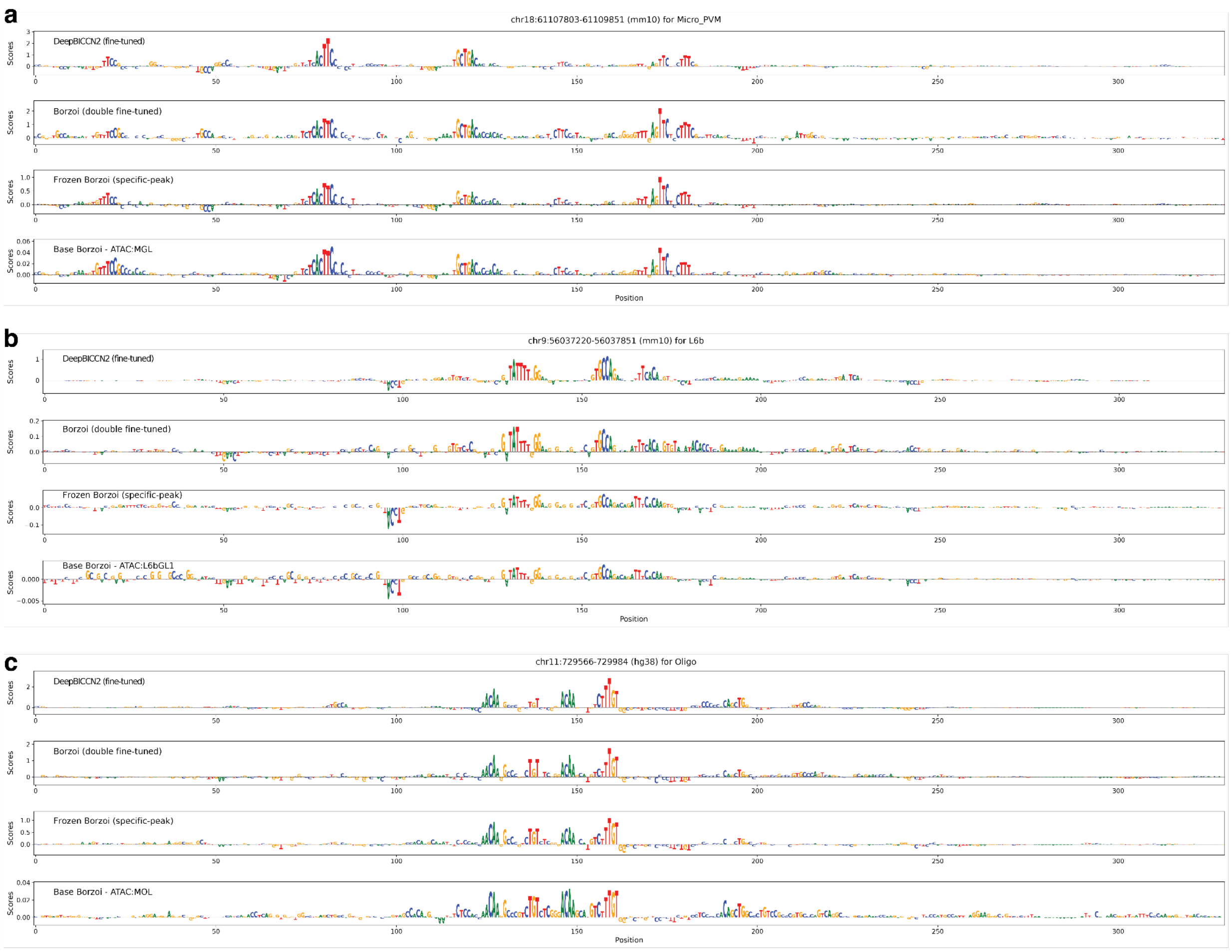


### Figure S16. Transfer learning model contribution scores for additional validated enhancers

DeepBICCN2, transfer learned Borzoi, and base Borzoi contribution scores for validated enhancers, calculated using expected integrated gradients. **(a)** Validated microglia enhancer[29] **(b-c)** Validated enhancers[23] from Fig. 2f not already shown in Fig. 5e.

#### Supplementary tables

##### Table S1. Synthetic zebrafish enhancers.

< *see TableS1.xlsx* >

Table S2. Key resource table.

< *see TableS2.xlsx* >
